# Occurrence and temporal dynamics of denitrifying protist endosymbionts in the wastewater microbiome

**DOI:** 10.1101/2025.07.24.665138

**Authors:** Louison Nicolas-Asselineau, Daan R. Speth, Linus M. Zeller, Ben J. Woodcroft, Caitlin M. Singleton, Lei Liu, Morten K. D. Dueholm, Jana Milucka

## Abstract

Effective wastewater treatment is of critical importance for preserving public health and protecting natural environments. Key processes in wastewater treatment, such as denitrification, are performed by a diverse community of prokaryotic and eukaryotic microbes. However, the diversity of the microbiome and the potential role of the different microbial taxa in some wastewater treatment plant setups is not fully understood. We aimed to investigate the presence and diversity of denitrifying bacteria of the candidate family *Azoamicaceae* that form obligate symbioses with protists in wastewater treatment plants. Our analyses showed that denitrifying endosymbionts belonging to the *Ca*. Azoamicus genus are present in 20-50% of wastewater treatment plants worldwide. Time-resolved amplicon data from four Danish wastewater treatment plants showed high temporal fluctuations in the abundance and composition of the denitrifying endosymbiont community. Fifteen high-quality metagenome-assembled genomes of denitrifying endosymbionts, four of which were circular, were recovered. Genome annotation showed that a newly described, globally widespread species, *Ca*. Azoamicus parvus, lacked a nitrous oxide reductase, suggesting that its denitrification pathway is incomplete. This observation further expands the diversity of metabolic potentials found in denitrifying endosymbionts and indicates that the contribution of microbial eukaryote holobionts to the wastewater ecosystem dynamics of nitrogen removal and greenhouse gas production may have been underestimated and needs further study.

## Introduction

Wastewater treatment plants (WWTPs) remove pollutants generated by agriculture, industry and domestic households from wastewater, and thus play a key role in protecting natural environments and health [1]. A taxonomically and metabolically complex microbial community carries out the chemical transformations inside WWTPs [2]. Based on previously published methods [3,4], an estimated 10^23^ bacterial individuals corresponding to 10^9^ species constitute the global activated sludge microbiome [5]. Only a few thousands of these species are common and occur in abundances relevant for the treatment process [6], and most of these species remain undescribed [7]. To date, most large-scale studies have focused on the diversity and role of prokaryotes (e.g. [5,6,8]). In contrast, the involvement of microbial eukaryotes, and particularly that of protists, remains understudied.

Protists, a paraphyletic group of eukaryotes with a unicellular level of organisation, are ubiquitous on Earth and inhabit diverse habitats, such as soil, freshwater and the ocean [9]. In WWTPs, protists have a documented function in shaping the prokaryotic community, mainly through predation [10]. However, protists also frequently form beneficial symbioses with bacteria and archaea [11]. These symbionts can fulfil a variety of roles for their hosts, including complementing metabolic functions [12], providing nutrition [13], or contributing to their competitive advantage [14] and motility [15].

A novel type of protist symbiosis was discovered recently, in which an obligate intracellular symbiont, *Candidatus* Azoamicus ciliaticola, generates ATP for its ciliate host through complete denitrification [16], the stepwise reduction of nitrate to dinitrogen via nitrite, nitric oxide and nitrous oxide. As such, *Ca*. A. ciliaticola fulfils a respiratory function strongly reminiscent of that of mitochondria, eukaryotic organelles dedicated to respiration and energy production. Since its original description in the anoxic hypolimnion of a stratified freshwater lake, relatives of *Ca.* A. ciliaticola have also been detected in groundwater environments on multiple continents [17]. The new groundwater species were identified based on the recovery of four circular metagenome-assembled genomes (MAGs). They form a monophyletic clade with *Ca.* A. ciliaticola but belong to two genera, *Ca.* Azoamicus and *Ca.* Azosocius, classified within the *Azoamicaceae* family in the UBA6186/Azoamicales order. Unlike the original lake symbiont detected in permanently anoxic waters, the genomes of all groundwater endosymbionts (with a size range between 284-374 kbp) encode a cytochrome-*cbb_3_* oxidase. This high affinity terminal oxidase [18] may allow the groundwater symbionts to perform aerobic respiration in addition to denitrification. The detection of *Ca*. Azoamicus aquiferis in oxic groundwater (6 mg/L dissolved oxygen) in the Hainich National Park expands the possible ecological niche of this symbiosis in comparison to the initial hypothesis [17]. Indeed, analyses of published 16S rRNA gene amplicons revealed that denitrifying endosymbionts are globally distributed, and their sequences can be found also in wastewater/activated sludge [17].

Denitrification is a key microbial process used in wastewater treatment for nitrogen removal (reviewed by [19]). Whereas most municipal biological WWTPs are designed to remove carbon in a single bioreactor with little nitrogen removal capacity, carbon removal is followed by dedicated nitrogen removal bioreactors in some WWTPs. Here, nitrogen, mainly found as ammonium, is removed from wastewater. In many of these nitrogen removal systems, a combination of nitrification/denitrification processes is employed, where ammonium and organic nitrogen compounds are oxidized to nitrate, which is then reduced via denitrification to mainly dinitrogen gas, a harmless product released into the atmosphere. Assimilation is another microbial process that may be responsible for some nitrogen and phosphorus removal. Due to imbalances between biological nitrous oxide (N_2_O) production and consumption, e.g. caused by incomplete denitrification, wastewater treatment processes may result in a net emission of N_2_O. Although WWTPs can generate substantial amounts of N_2_O, they remain a minor contributor to anthropogenic N_2_O emissions [20].

This study aimed to comprehensively investigate the ecology of denitrifying endosymbionts in WWTPs using molecular and imaging techniques, expanding on the initial findings of [17]. Specifically, we explored the diversity of denitrifying endosymbionts and their hosts in WWTPs at a genomic and genetic scale, and retrieved novel MAGs that provide new insights into their metabolic potential. We additionally assessed their prevalence in global WWTP datasets. Finally, we evaluated the temporal variations in abundance and composition of the individual endosymbiont lineages in four municipal WWTPs in Denmark.

## Materials and Methods

### Recovery of denitrifying endosymbiont genomes

MAGs of denitrifying endosymbionts were recovered through three approaches. First, metagenomic assemblies obtained from activated sludge samples by [21] were aligned with the complete genomes of already known denitrifying symbionts, i.e. *Ca*. A. ciliaticola and its four relatives from groundwater [17], using BLASTN v. 2.6.0 [22]. Aligning contigs > 3 kb were extracted to form a MAG of a denitrifying symbiont.

Secondly, publicly available metagenomes present in the Sandpiper v. 0.0.23 [23] database were searched for the presence of denitrifying symbiont sequences. This database was built following the SingleM pipeline [23]. Briefly, metagenomes were screened for reads covering conserved regions of single-copy marker genes when translated to amino acids. The corresponding nucleotide sequences were clustered into operational taxonomic units subsequently assigned with a taxonomy. As a result, taxonomic profiles are available for all metagenomes found in Sandpiper. Around 6000 published metagenomes that contained at least one operational taxonomic unit classified within the UBA6186 order were picked. This number was reduced to 4000 by keeping only datasets that hit translated sequences of the *Ca.* A. ciliaticola and four groundwater related closed genomes. Among them, 421 metagenomes sequenced from wastewater systems and displaying a coverage above 5 for denitrifying endosymbiont sequences were downloaded with fasterq-dump v. 2.11.0 (https://github.com/ncbi/sra-tools). Additionally, the 227 metagenomes from WWTPs published by the Global Water Microbiome Consortium (GWMC, [5]) were downloaded with fasterq-dump and added to the collection of datasets to trawl for denitrifying endosymbiont presence.

Raw reads from the selected datasets were quality-filtered to keep a Phred score above 20 and adapter trimmed using TrimGalore v. 0.6.7 (https://github.com/FelixKrueger/TrimGalore). To limit the amount of metagenomic datasets to assemble, we trawled them using a read-based pre-screening approach to further narrow down the selection from Sandpiper and GWMC. The datasets were screened for denitrifying endosymbiont genomes by sequentially looking for marker genes, first for the presence of the *tlcA* gene coding for the ATP/ADP translocase, with remaining datasets screened for the *nosZ* gene coding for the nitrous oxide reductase. The BLAST Score Ratio [24,25] was used for this approach, as described in the Supplementary Methods.

The 27 metagenomes from Sandpiper and GWMC selected based on the *tlcA* and *nosZ* BLAST Score Ratio (Table S4) were assembled with MEGAHIT v. 1.2.9 [26] applying the meta-large presets parameter. Similarly to the data from [21], the resulting assemblies were aligned with the five known complete genomes of denitrifying endosymbionts with BLASTN v. 2.6.0. Aligning contigs > 3 kb from each dataset were binned into a MAG.

The obtained MAGs were further refined as described in the Supplementary Methods. A MAG was considered complete when the assembly graph provided by SPAdes [27] and visualised in Bandage [28] was closed and circular. A visualisation of gene clusters among the published and wastewater MAGs of denitrifying endosymbionts was generated using clinker v. 0.0.28 [29] to identify contigs that may result from binning error.

Lastly, seven denitrifying endosymbiont MAGs were retrieved from the Microbial Database of Activated Sludge and Anaerobic Digesters (MiDAS, [6,8]) global genome catalogue (accession PRJEB83983). It is based on Nanopore long-read sequencing, assembly and binning of representative samples from the MiDAS global survey of WWTPs [6], as detailed in the Supplementary Methods.

All MAGs were classified as high-quality or medium-quality MAGs based on the criteria defined by [30] that include completion, contamination and assembly statistics such as N50, largest contig and number of contigs. As the recommended tools for MAG completeness assessment are not designed for tiny genomes, we could only estimate this parameter (Table S5), and mainly based the quality sorting on the other criteria defined by [30]. All recovered MAGs were annotated as described in the Supplementary Methods.

### Phylogenomic analyses

A sequence alignment including the wastewater MAGs, the other known complete denitrifying endosymbiont genomes and four UBA6186 sister clade genomes retrieved from GTDB (GCA_002422785, GCA_903822055, GCA_903827925, GCA_903907935; [31]) was built in anvi’o v. 7.1 [32]. An anvi’o contigs database was created for each genome. The amino acid sequences of 71 bacterial single-copy marker genes (list of genes from [33], 13001 amino acids) identified by anvi’o in each MAG using HMMER (http://hmmer.org/) were aligned through the MUSCLE v. 3.8.1551 aligner tool with default parameters [34] and concatenated. A phylogenomic tree was generated based on this alignment by IQ-TREE v. 2.2.6 [35] using ModelFinder [36] for model selection (best-fit: mtInv+F+I+R4) and UFBoot2 [37] for ultrafast bootstrap approximation (1000 replicates), and displayed in iTOL [38].

### Wastewater sampling and metagenome/metatranscriptome sequencing

Wastewater was sampled from the aeration and anoxic tanks of the *hanseWasser* WWTP (53.115 N 8.7105 E, hereafter Seehausen WWTP) in Bremen, Germany, in October 2022. The aeration tank was sampled again in August and November 2023. This WWTP operates following the anaerobic-anoxic-oxic process. Wastewater was collected into Duran bottles, kept at ambient temperature and processed within an hour. Further information about the sampling and processing methods of the samples can be found in Table S1. Total DNA and RNA were extracted for metagenome and metatranscriptome sequencing, as described in the Supplementary Methods.

The presence of denitrifying endosymbionts in the Seehausen WWTP was determined by mapping the metagenomic reads onto the wastewater MAGs. A genome was considered present in a metagenome when the calculated breadth of coverage was within 15% of the expected breadth according to the formula: expected breadth = 1 – e^(−0.883*depth of coverage) [39,40]. More details can be found in the Supplementary Methods.

<u>Double labelling of oligonucleotide probes for fluorescence *in situ* hybridisation (FISH)</u> Wastewater samples were left undisturbed for up to two hours after sampling until the solid fraction had settled. Ten mL of the relatively clear supernatant were fixed with 2% paraformaldehyde (Electron Microscopy Sciences, USA) for 1-2h at room temperature. Fixed samples were filtered onto 3 µm-Isopore^TM^ polycarbonate filters (Merck Millipore, USA) using gravity filtration.

Filters were dried and embedded in 0.2% MetaPhor^TM^ Agarose (Lonza, Switzerland) and first hybridised with the oligonucleotide probe plagi1083 (5’-TTGTGTCCATACTTCCCCC-3’) double labelled with Atto594 for 2h at 35% formamide and 46°C. The plagi1083 probe design and evaluation for FISH are described in the Supplementary Methods. The same filter pieces were additionally hybridised with the oligonucleotide probe eub62A3_813 (5′-CTAACAGCAAGTTTTCATCGTTTA-3′) double labelled with Atto488 dye (Biomers, Germany) for 3 to 5h at 25% formamide concentration and 46°C and washed as described in [16]. The eub62A3_813 probe targets all the denitrifying endosymbionts detected in the Seehausen WWTP, with the notable exception of *Ca.* A. parvus. The filters were further stained with 1 µg/mL 4′,6-diamidino-2-phenylindole (DAPI) for 10 minutes at 4°C in the dark, then washed twice in milliQ water. Dried filters were embedded in 4:1 Citifluor Vectashield (Vector Laboratories, USA) and image stacks were recorded on a confocal laser scanning microscope (Zeiss LSM 780, Germany; 63× oil objective, 1.4 numerical aperture, maximum intensity projection). As a negative control, filters were labelled with the NON388 probe (5’-ACTCCTACGGGAGGCAGC-3’) and imaged at the same respective laser intensities. Because the wastewater samples showed autofluorescence in the endosymbiont FISH channel (Fig. S1), we made sure that the fluorescence in the endosymbiont channel always overlapped with a DAPI signal and that these signals were absent in the negative control. The used probes are summarised in Table S2.

### SSU rRNA phylogenetic analyses

Two full-length 18S rRNA gene sequences putatively belonging to denitrifying endosymbiont hosts were recovered from a bioreactor inoculated with wastewater in Michigan and from the Seehausen WWTP, as described in the Supplementary Methods. The Seehausen sequence was aligned with 18S rRNA gene sequences used in the host phylogenetic tree from [16] with the addition of the full-length host sequences extracted from groundwater [17] using MUSCLE v. 5.1 with default parameters. A phylogenetic tree was computed with IQ-TREE v. 2.2.6 using ModelFinder (best-fit: TIM3+F+R3) and UFBoot2 for ultrafast bootstrap approximation (1000 replicates) and displayed in iTOL. Another phylogenetic tree that includes the 18S rRNA gene sequence from the Michigan samples was computed in the same way (best-fit model: TN+F+I+R3).

For 16S rRNA gene phylogenetic investigations, the gene sequences of all members of the midas_f_1324 family were extracted from the MiDAS v. 5.3. The 16S rRNA gene sequences of the wastewater MAGs were associated with a MiDAS taxonomy using the best BLAST hit. Species belonging to the midas_f_1324 family, which were not detected in amplicons from the sampled WWTPs were removed from the tree in Fig. 2b but can be found in Fig. S2. The alignment and maximum likelihood phylogenetic tree were computed with a similar method as for the 18S rRNA gene-based tree (ModelFinder best-fit: TVM+F+R3).

### Assigning a MiDAS taxonomy to 16S rRNA gene amplicon datasets

The 16S rRNA gene sequences of denitrifying endosymbionts previously recovered from freshwater lake, groundwater and wastewater were blasted against the MiDAS v. 5.3 reference database. All denitrifying endosymbionts hit reference sequences belonging to the midas_f_1324 family, which was not the case of the non-denitrifying symbiont species in the UBA6186 order. The midas_f_1324 family was thus used as an approximation of the *Azoamicaceae* family.

All 16S rRNA gene amplicons published by MiDAS 4 and GWMC as well as time series datasets from [38] were trawled for amplicon sequencing variants (ASVs) belonging to the midas_f_1324 family. Briefly, the amplicon processing in QIIME 2 v. 2022.11.1 [39] included read trimming, quality filtering, clustering into ASVs with > 99% identity and MiDAS v. 5.3 taxonomy classification. Further details on the amplicon treatment can be found in the Supplementary Methods.

## Results

### Recovery of denitrifying endosymbiont MAGs with diverse potentials for energy production

To assess the diversity of denitrifying endosymbionts in WWTPs, we retrieved a total of 16 MAGs from sequencing datasets assigned as wastewater (Fig. 1, Table S4). Six MAGs were recovered from our own screening of public datasets (see Methods), three from metagenomic assemblies from 23 Danish WWTPs reported by [21], and the other seven from the genome catalogue obtained by long-read sequencing of 83 samples from the MiDAS global survey of WWTPs [6].

**Figure 1:**
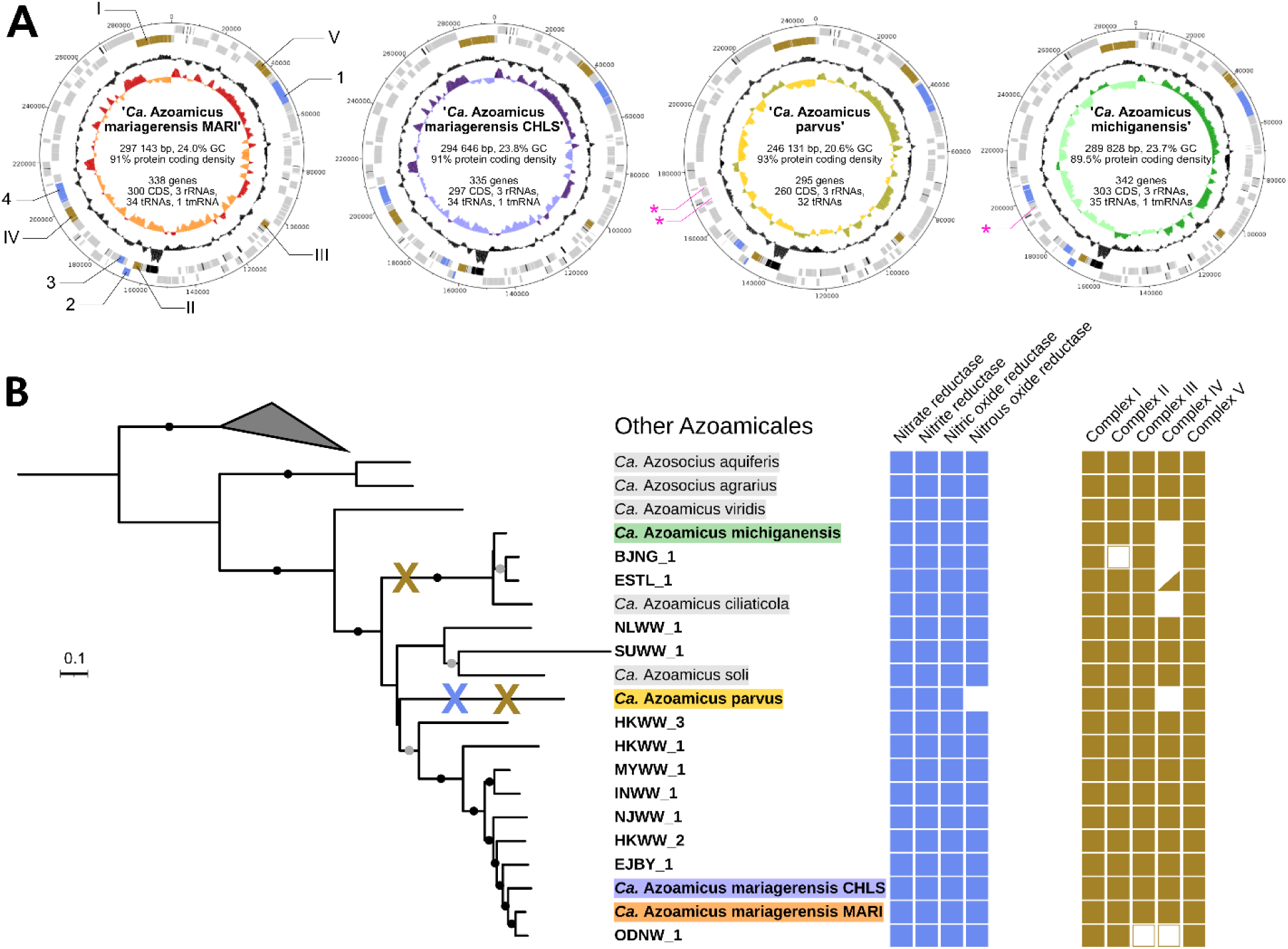
Genomic diversity of denitrifying endosymbiont MAGs. A, Circular DNA plots and characteristics of complete genomes of denitrifying endosymbionts. For each plot, rings from the outside to the inside correspond to forward features, reverse features, GC content and GC skew. Coding sequences are shown in grey, rRNA and tRNA in black, coding sequences involved in denitrification and electron transport chain in blue and brown respectively. For GC content and GC skew plots, lighter areas reflect below-average values and darker areas above-average values. When present, the position of highlighted genes is conserved in all four genomes. For visual clarity, the genes are only annotated in the *Ca.* A. mariagerensis MARI genome and pink asterisks denote the loci of lost genes. 1: *narGHIJKT*; 2: *nirQ*, *norD*; 3: *nirK*, *norBC*; 4: *nosDLFRYZ*; I: *nuoABCDEFGHIJKLMN*; II: *sdhABCD*; III: *qcrABC*; IV: *ccoNOPQS*; V: *atpABCDEFGHI*. **B,** Phylogenomic tree of all wastewater MAGs and other known denitrifying endosymbionts and associated genetic potential for energy production. MAGs retrieved from wastewater are shown in bold and complete genomes have a highlighted background matching the colours used in panel A. Genomes from previous works are highlighted in grey. Black and grey dots on the branches denote bootstrap values above 90% and 80% respectively. An enzyme was considered encoded (filled square) when all genes coding for functional core subunits were present in the genome, possibly encoded (empty square) when the genes were absent in an incomplete genome but suspected to be missing from the assembly, and absent (no square) when no gene was found at the expected locus on a contig based on synteny. The triangle indicates the presence of a cytochrome-*cbb_3_* oxidase gene operon of *Azoamicaceae* origin in the ESTL_1 MAG, but its contig contains no other gene, making its association with the MAG uncertain. Blue and brown crosses on the tree indicate the loss of nitrous oxide reductase and cytochrome-*cbb_3_* oxidase respectively.

**Figure 2:**
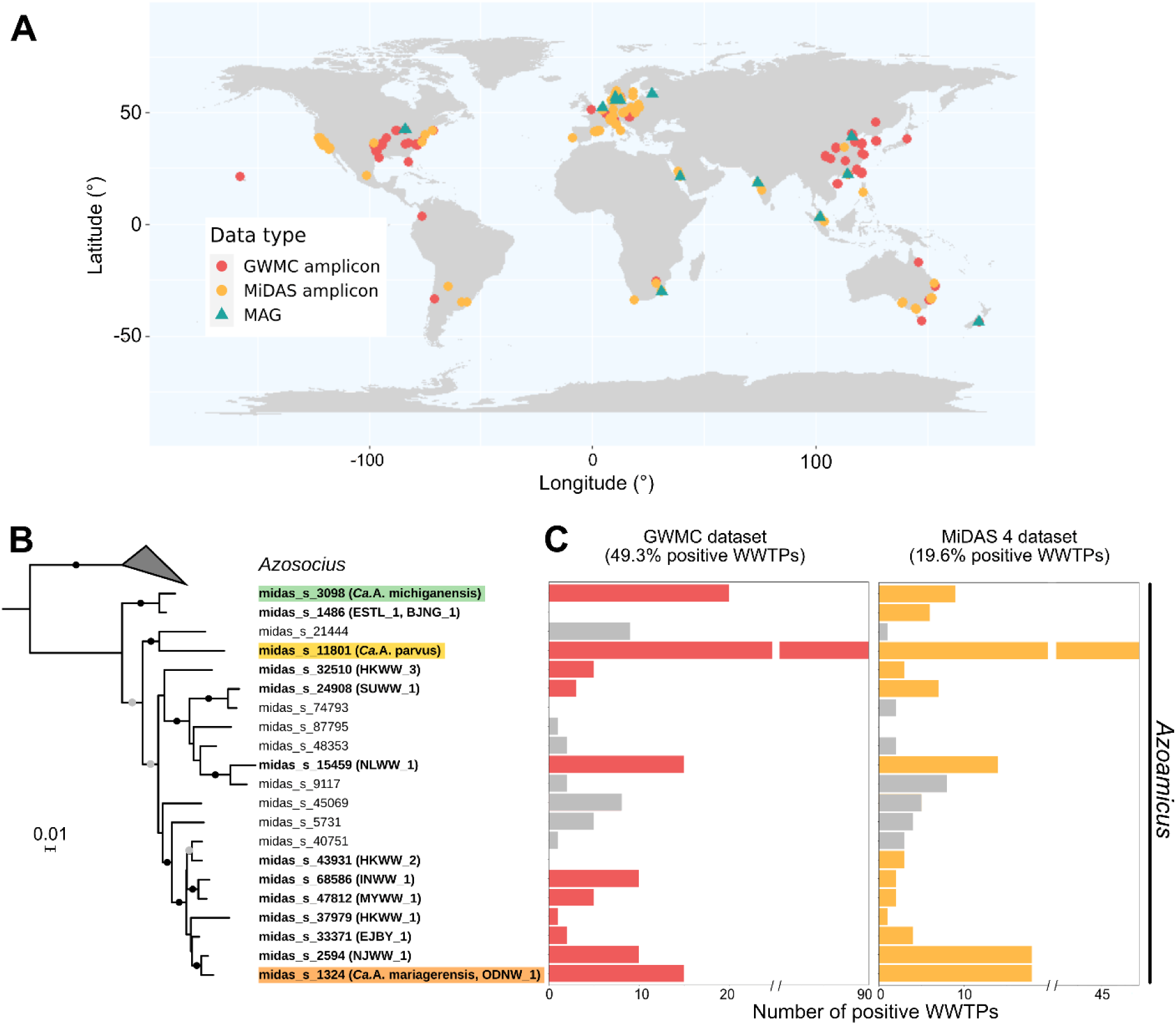
Occurrence and phylogeny of denitrifying endosymbiont species in WWTPs across the world. A, Location of WWTPs in which *Azoamicaceae* endosymbionts were identified either through 16S rRNA gene amplicon sequencing or through the retrieval of whole genomes. Colour indicates the source dataset. **B,** 16S rRNA gene sequence-based maximum likelihood phylogenetic tree of MiDAS 5.3 species presumably belonging to the *Azoamicaceae* family. Species to which the wastewater MAGs belong are shown in bold. Black and grey dots on the branches denote bootstrap values above 90% and 80% respectively. **C,** Number of WWTPs sampled as part of GWMC and MiDAS 4 in which each MiDAS 5.3 species is present. Lineages for which no MAG is available are represented in grey.

Overall, 15 of the 16 recovered MAGs were estimated to be over 90% complete and included 23S, 16S and 5S rRNA genes. They were thus considered high-quality MAGs [30]. One MAG (BJNG_1; 234 kbp, 24 contigs) was medium-quality because it lacked a full-length 16S rRNA gene (Table S5). The overall length of the MAGs varied between 234 and 303 kbp. Notably, four of the high-quality MAGs were circular and complete (Fig. 1a). Based on their genome characteristics and synteny with previously published *Azoamicus* genomes [16,17] (Fig. S3), we propose that they belong to three new species within the *Azoamicus* genus of the *Candidatus* Azoamicaceae family (Supplementary Discussion, Table S6). We name them *Candidatus* Azoamicus mariagerensis strains MARI and CHLS, *Candidatus* Azoamicus parvus and *Candidatus* Azoamicus michiganensis (see Etymology).

All key metabolic pathways previously described in denitrifying endosymbionts were encoded in each MAG, supporting a role in the production and exchange of energy with the host (Fig. 1b, Fig. S4). A complete denitrification pathway including a nitrate reductase (*narGHI*), a nitrite reductase (*nirK*), a nitric oxide reductase (*norBC*) and a nitrous oxide reductase (*nosZ*) was encoded in nearly all MAGs (Supplementary Discussion). In contrast to other known genomes, the complete genome of *Ca*. A. parvus lacked the *nosZ* gene (Fig. 1b, Table S7). A closer search showed that in fact the complete operon (consisting of *nosDLFRYZ*) was missing. As *Ca*. A. parvus still encodes the rest of the denitrification pathway, i.e. NarGHI, NirK, NorBC, it appears to reduce nitrate to nitrous oxide but not further to dinitrogen gas. The core complexes of the electron transport chain (complexes I, II, III and V) were encoded in all MAGs, specifically the NADH dehydrogenase complex (*nuoA* to *nuoN* genes, complex I), the succinate dehydrogenase (*sdhABCD*, complex II), the cytochrome *bc*_1_ (*qcrABC*, complex III) and the ATP synthase (*atpA* to *atpH*, complex V; Fig. 1b). Only two MAGs, BJNG_1 and ODNW_1, lacked genes coding for some catalytic subunits of complexes II and III respectively. However, this is likely due to MAG incompleteness as the complex II and III genes in those two MAGs were either located on the edge of a contig or presumed to be found in between two contigs based on gene synteny.

As observed in the four recently described groundwater species [17], most wastewater MAGs (12 of 16; Fig. 1b) possessed a complete operon encoding a cytochrome-*cbb*_3_ oxidase (*ccoNOPQ*, complex IV) and thus the potential to respire oxygen in addition to nitrate. The operon was not found in the genomes of *Ca.* A. michiganensis, *Ca*. A. parvus, BJNG_1 and ESTL_1 (Supplementary Discussion). Finally, an ATP/ADP translocase (*tlcA*), a presumed key protein for the ATP-providing function of the symbiont, was present in all MAGs. Thus, even with an increasing taxonomic diversity, the respiratory functions of denitrifying symbionts remain preserved in species from various habitats.

Phylogenomic analyses (Fig. 1b) revealed that the wastewater symbiont species were not monophyletic. While one clade within the *Azoamicaceae* consisted exclusively of genomes retrieved from wastewater environments, two other clades containing wastewater-derived lineages also included a groundwater and a lacustrine species. *Ca*. A. michiganensis, BJNG_1 and ESTL_1 formed a distinct clade with freshwater species *Ca*. A. ciliaticola, while NLWW_1 and SUWW_1 branched with *Ca*. A. soli described from groundwater.

### Global distribution of denitrifying endosymbionts

We used datasets from the GWMC [5] and MiDAS 4 [6] sampling campaigns to estimate the geographical distribution and occurrence of denitrifying endosymbionts worldwide. These datasets consist of 16S rRNA gene amplicons sequenced from WWTP samples around the world (activated sludge for GWMC, mostly activated sludge and some anaerobic digesters for MiDAS 4). The MiDAS project was of particular interest, as it provides a consistent ecosystem-specific reference taxonomy, MiDAS v. 5.3, for microbial taxa detected in wastewater to the species resolution. Due to differences between genome and 16S rRNA MiDAS 5.3 taxonomies, we first established that the *Azoamicaceae* all belonged to the midas_f_1324 family by linking the 16S rRNA genes from the symbiont MAGs to the 16S rRNA genes in MiDAS 5.3 reference taxonomy. This allowed us to leverage the MiDAS 5.3 taxonomic framework for the detection of denitrifying endosymbiont species in both the MiDAS 4 and GWMC amplicon datasets.

Denitrifying endosymbionts putatively belonging to the *Azoamicaceae* (midas_f_1324) family were found on all inhabited continents (Fig. 2a), despite the fact that some regions, e.g. the African continent, are strongly undersampled by both MiDAS and GWMC (Fig. S5). 16S rRNA gene sequences available in the MiDAS v. 5.3 dataset revealed the existence of some *Azoamicaceae* species for which a high-quality MAG is still missing (Fig. 2b). However, these lineages were generally less frequently detected in the screened datasets (Fig. 2c), indicating that they are less globally distributed and abundant than the taxa with recovered MAGs. Overall, the 16S rRNA gene-based taxonomy (Fig. 2b) was largely congruent with the concatenated marker gene phylogeny (Fig. 1b). A notable exception was *Ca*. A. parvus, which branches deeper in the 16S rRNA gene tree, with a higher bootstrap support.

*Azoamicaceae* were detected in 49.3% (132 out of 268) of the individual WWTPs sampled by the GWMC and in 19.6% (123 out of 626) of the ones from MiDAS 4 (Fig. 2c). Denitrifying endosymbionts and their hosts seemed generally successful and widespread in activated sludge and could thus be considered prevalent members of its microbiome. The large difference between the two detection values might be explained by variations in the types of sample (activated sludge or anaerobic digester) and in amplification protocols. *Ca.* A. parvus was particularly widespread compared to other species and present on a global scale (Fig. 2c). In general, *Azoamicaceae* symbionts were found in multiple types of WWTPs treating a range of wastewater strength and operating with a large variety of conditions (Fig. S6), suggesting that they can cope with a range of different conditions.

### Temporal variations of denitrifying endosymbiont abundance

To check whether the lack of detection of *Azoamicaceae* in certain WWTPs could be due to temporal fluctuations, we assessed the abundances of the different *Azoamicaceae* species in four Danish WWTPs (Aalborg East and West, Damhusåen and Randers) over ca. 2000 days. These four plants have the same design and operate with similar conditions, e.g. 4-5g/L of suspended solids and 0.5-2mg/L of dissolved oxygen. We quantified the reads belonging to *Azoamicaceae* in the high-resolution time-series 16S rRNA gene amplicon datasets. This analysis showed that denitrifying endosymbiont abundances dramatically varied over time (Fig. 3). Occasionally, their relative abundance spiked suggesting the occurrence of blooms of denitrifying endosymbionts, such as the sudden short-lived occurrence of midas_s_11801 (*Ca.* A. parvus) in Damhusåen WWTP around days 1200-1300 of the time series. Similar fluctuations are also observed in the symbiont read counts (Fig. S7). Some denitrifying endosymbiont species appeared in specific WWTPs seasonally, e.g. midas_s_5731 was typically detected in the Randers WWTP in summer and midas_s_1324 (*Ca*. A. mariagerensis) present around the fall period in the Aalborg East WWTP. This suggests that the local *Azoamicaceae* populations were influenced by the origin and conditions of wastewater production, as previously observed for prokaryotes across various industrial [41] and municipal WWTPs [42]. As these four plants all treat a mix of municipal wastewater and industrial discharge, the distribution of wastewater types, the production location as well as the housing characteristics could explain some patterns of occurrence of specific *Azomicaceae* species in individual WWTPs.

**Figure 3:**
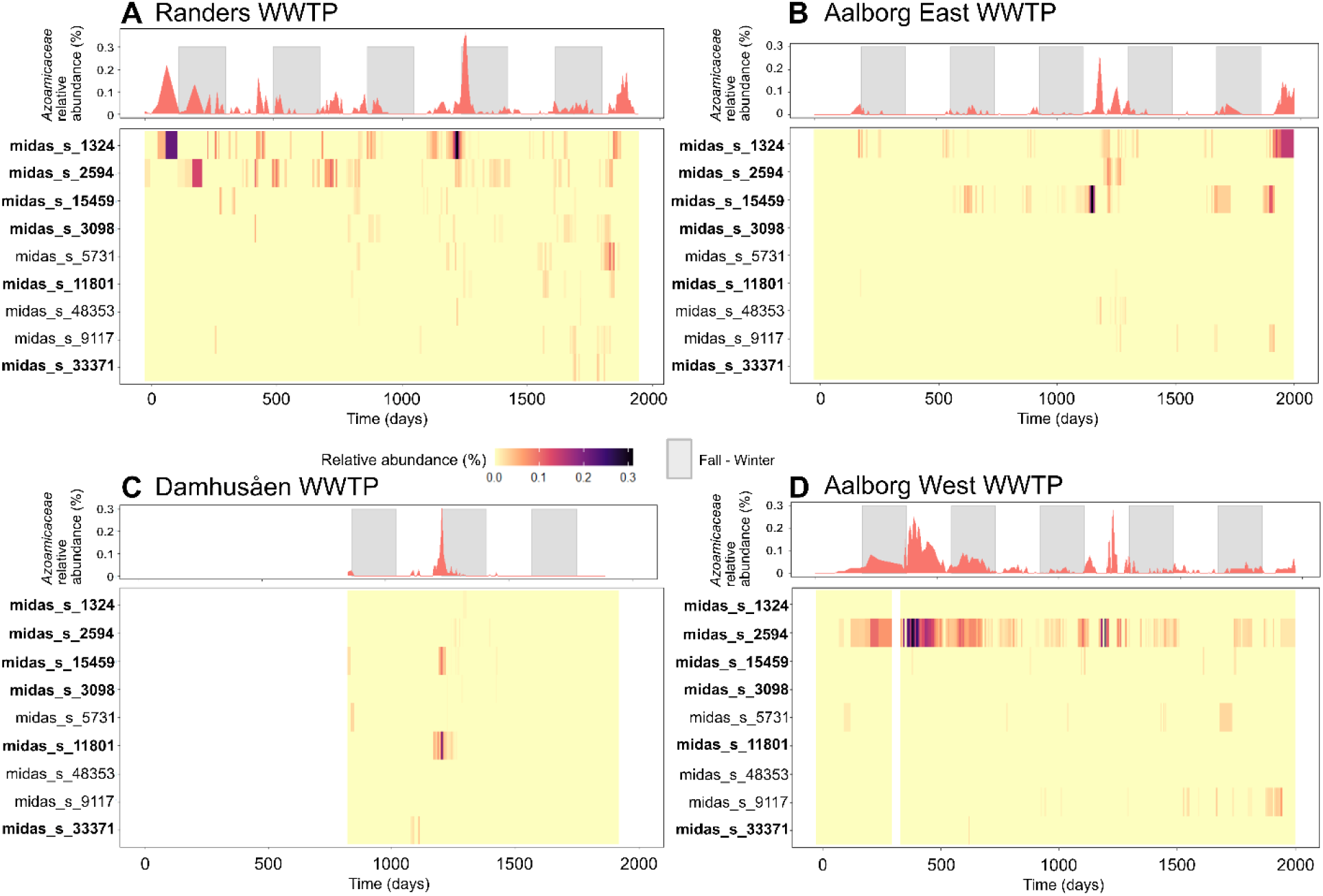
Relative abundance of denitrifying endosymbionts over time in four Danish WWTPs. Each panel depicts the community fluctuations at the family and species level in the **A,** Randers, **B,** Aalborg East, **C,** Damhusåen, **D,** Aalborg West WWTPs. All listed MiDAS 5.3 species are expected to belong to the *Azoamicaceae* family. A MAG is available for the species in bold. Grey bars on the upper plots highlight fall-winter time defined as the period between 21 September and 21 March.

Interestingly, also the composition of the denitrifying endosymbiont communities varied with time. In all four investigated WWTPs, multiple symbiont species were observed to coexist, albeit to a different degree. For example, in the Randers WWTP up to seven symbiont species were detected simultaneously at roughly comparable abundances and all contributed to the observed spikes in total *Azoamicaceae* abundances. On the contrary, in the Aalborg West WWTP, one species, midas_s_2594 (NJWW_1), strongly dominated the denitrifying endosymbiont community.

Overall, in the four Danish WWTPs, *Azoamicaceae* could be detected in 26-63% of the sampling points (Supplementary Discussion). While denitrifying endosymbionts are widespread in WWTPs worldwide, their abundance and diversity can be expected to strongly fluctuate over time. These organisms may thus be present in more WWTPs worldwide than suggested based on the available amplicon data.

### Putative host association of denitrifying endosymbionts in WWTPs

The newly revealed diversity of denitrifying symbionts in wastewater ecosystems raises the question of the host identity. So far, lacustrine and groundwater symbiont lineages have been shown and proposed, respectively, to always associate with ciliates of the Plagiopylea class [16,17]. Therefore, we tried to recover 18S rRNA gene sequences in short-read metagenomes from which symbiont MAGs were successfully retrieved (Table S4). Unfortunately, only from one of them, a bioreactor sample from Michigan, USA, could a full-length 18S rRNA gene sequence be recovered (Fig. S8). This sequence was very closely related to the 18S rRNA gene belonging to the host of the lacustrine *Ca*. A. ciliaticola (97.8% identity). As this dataset contained the highest abundance of one of the wastewater symbiont species, *Ca*. A. michiganensis, it is likely that the unsuccessful recovery of the host 18S rRNA sequence in other datasets is due to the low abundance of the putative host.

Therefore, to further investigate the host identity of the wastewater symbiont lineages, we collected samples from our local municipal Seehausen WWTP (Bremen, Germany). Three sampling campaigns were performed over the course of one year, during which the supernatant and sludge of the aeration and anoxic reactors were collected. In all samples, multiple species of denitrifying symbionts were found, with *Ca.* A. mariagerensis often being the most abundant (Fig. 4a, Fig. S9; Supplementary Discussion). Of the three sampling campaigns, the abundance of denitrifying symbionts was lowest in August 2023 and relatively high in October 2022 and November 2023 (Fig. 4a), which is also reflected in the gene transcription levels (Fig. S10; Supplementary Discussion). From the October 2022 metagenome, a full-length 18S rRNA gene sequence belonging to a Plagiopylean ciliate and closely related to the *Ca.* A. ciliaticola and *Ca.* A. michiganensis host sequences was assembled (Fig. 4b). This 18S rRNA sequence was most closely related to a ciliate sequence retrieved from a WWTP in Sendai, Miyagi, Japan. Due to the high abundance of *Ca.* A. mariagerensis MARI in the Bremen WWTP at the time of sampling, we speculate that the retrieved 18S rRNA gene sequence could belong to its host.

**Figure 4:**
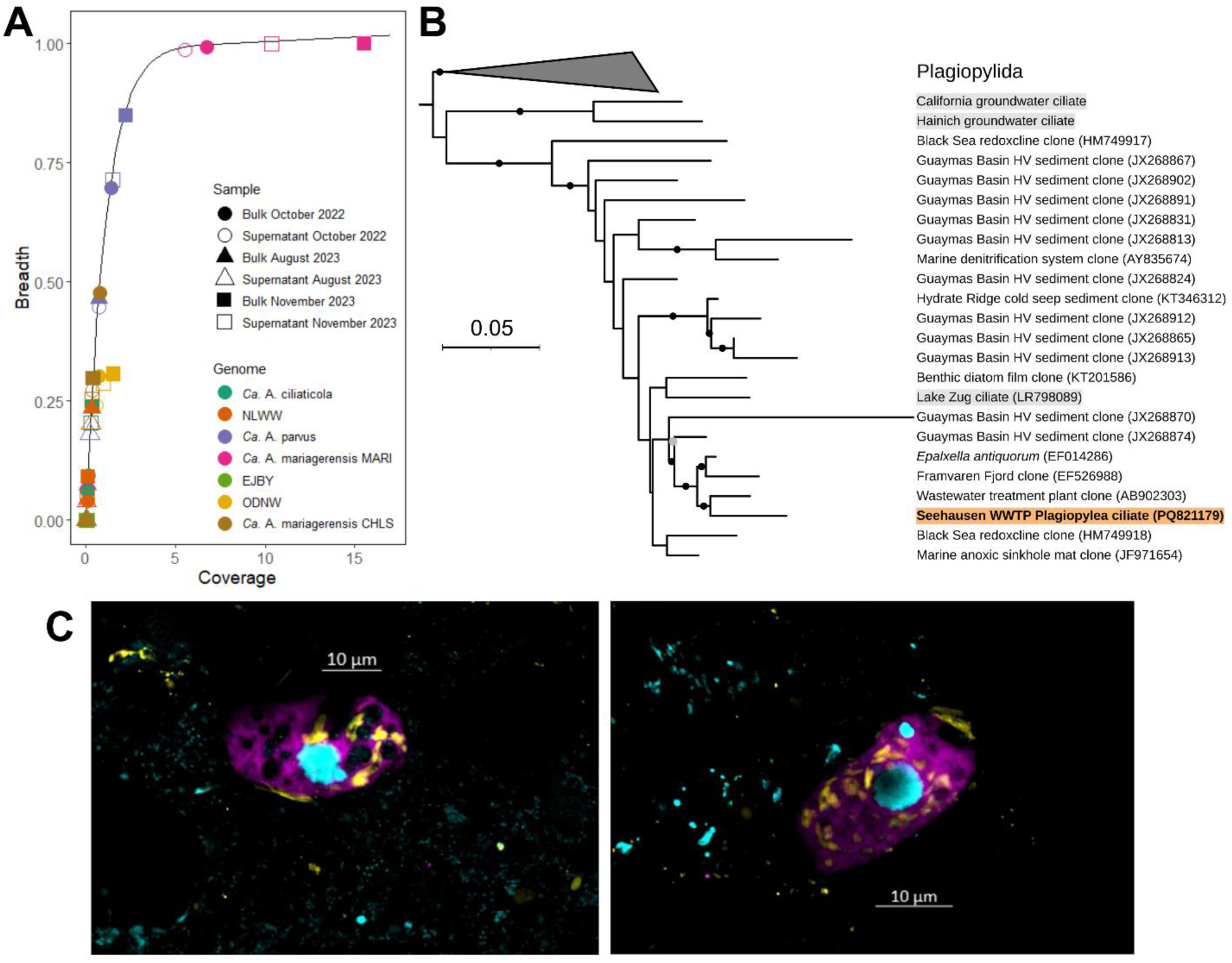
Diversity of denitrifying endosymbionts within the Seehausen WWTP. A, Detection and abundance of denitrifying endosymbiont lineages in fractions of wastewater sampled over time in the aeration tank. The black line corresponds to the expected breadth for each coverage value according to the formula described in the Methods. Symbols on the line thus indicate specific detection of symbiont MAGs. Coverage: mean number of reads aligning each base. Breadth: fraction of a genome covered by at least one read. **B,** 18S rRNA gene based maximum likelihood phylogenetic tree showing the sequence of the putative host of *Ca*. A. mariagerensis MARI from the Seehausen WWTP (orange background, bold font), as well as sequences of non-wastewater ciliates potentially associated with denitrifying endosymbionts (grey background). Black and grey dots on the branches denote bootstrap values above 90% and 80% respectively. **C**, Double labelling of oligonucleotide probes for FISH pictures of denitrifying endosymbionts and their ciliate host in the aeration tank in November 2023 imaged with confocal laser scanning microscopy. The Plagiopylean probe is shown in violet, the denitrifying endosymbiont specific probe in yellow and DAPI staining in blue. Images of the individual probes and negative controls can be found in Fig. S1.

To conclusively link the wastewater symbionts with their presumed Plagiopylean hosts, we designed a specific oligonucleotide probe (plagi1083 targeting the 18S rRNA gene) for the visualisation of the putative host class in a sample from the Seehausen WWTP from November 2023. Unicellular eukaryotes targeted by this probe were detected in all tested samples (Fig. 4c; Fig. S11), confirming the presence of the putative host lineage. The predominant morphotype included ca. 20 μm-long oval cells with a macro– and micro-nucleus (Fig. 4c). We then combined this probe with the previously used 16S rRNA gene probe (eub62A3_813, [16]) for simultaneous visualisation of the host and the symbiont. A positive signal for the 16S rRNA probe was solely detected inside the ciliates that hybridised with the Plagiopylea-specific 18S rRNA probe (Fig. 4c), confirming that the abundant wastewater lineage of denitrifying symbionts resides inside Plagiopylean ciliate hosts.

## Discussion

### Presence of denitrifying endosymbionts in WWTPs around the world

Denitrifying endosymbionts of anaerobic ciliates have only recently been discovered in freshwater lakes [16] and groundwater ecosystems [17]. A search of global 16S rRNA gene amplicons indicated that these organisms might also be prevalent in wastewater [17]. Indeed, our survey of global wastewater sequencing datasets revealed the presence of diverse species of *Ca*. Azoamicus symbionts in WWTPs worldwide. The success of *Azoamicaceae* in WWTPs shows that their ecological niche includes engineered systems that display strikingly different conditions to pristine natural environments. WWTPs typically have higher nutrient concentrations than the environments where the symbiosis was observed before, as well as high microbial diversity, competition and turnover [19]. While physicochemical conditions such as pH, temperature and oxygen concentration are monitored and stabilised [43], these parameters can vary greatly among the individual WWTPs.

The activated sludge microbiome is thought to be affected by both the environmental parameters [44] and the influent wastewater [45]. However, the activated sludge community composition is also shaped by stochastic factors [5], highlighting the difficulty of predicting dynamics shaping microbial communities in WWTPs. Denitrifying endosymbionts were detected in a broad range of WWTP conditions, with no apparent link between specific symbiont lineages and operating parameters (Fig. S6).

Among the four Danish WWTPs in which the microbiome was monitored over time, no general temporal pattern for the occurrence of *Azoamicaceae* could be determined. Some species of denitrifying endosymbionts exhibited seasonally recurring blooms, similar to other members of activated sludge communities [42,46]. However, most lineages appeared and disappeared seemingly randomly. Since only a limited portion of microorganisms entering WWTPs via influent water persist in the activated sludge [44], microbes that regularly immigrate into a WWTP have a higher chance to survive for a longer time inside the plant, and appear growing [47]. We speculate that some of the observed drops in the abundance of discrete *Azoamicaceae* lineages may be a result of their interrupted immigration into a WWTP.

Based on the assumption that the placeholder midas_f_1324 family approximates the monophyletic clade of *Azoamicaceae*, we could detect denitrifying endosymbionts in 20-50% of WWTPs worldwide. However, given the observed fluctuations in the symbiont community abundance and composition over time, sampling time is likely a major factor in their detection. Hence, the true distribution of these symbionts across WWTPs might be even broader.

### Features of denitrifying symbionts in WWTPs

The new symbiont genomes and host 18S rRNA sequences expand the known diversity of denitrifying symbioses, particularly in the *Ca.* Azoamicus genus that captures most of the species diversity of denitrifying endosymbionts. The topology of the symbiont phylogenomic tree and host 18S rRNA gene tree remained congruent (Figs. 1a and 4a), although the bootstrap values of the latter are sometimes low. *Ca.* Azosocius and *Ca.* Azoamicus symbionts seem to form distinct monophyletic clades, just like their respective hosts. This further corroborates the long-term vertical inheritance of denitrifying symbionts suggested by [17].

Interestingly, the wastewater species do not form a monophyletic clade. Given the young age of WWTPs, it is likely that the diversification of wastewater denitrifying endosymbionts and their hosts takes place in environments upstream of immigration into WWTPs. Since we could so far not detect the wastewater species in any natural habitat, the nature of these upstream environments remains to be determined.

All new wastewater species have a highly conserved genome structure and overall gene synteny (Fig. S3), corroborating the concatenated marker gene phylogeny (Fig. 1b). The genome sizes of the wastewater species range from 246 to 303 kbp, revealing that some genes have been selectively lost in distinct clades. Genome erosion is a well-documented mechanism in obligate bacterial endosymbionts of e.g. insects [48]. Inside the host cells, many endosymbiont genes become superfluous or redundant with the host genetic machinery and may get lost [49]. This is the case of genes related to e.g. replication, transcription and translation [16]. The extent of genome erosion broadly reflects the level of integration of symbionts inside the host cell, and the set of genes retained in tiny endosymbiont genomes typically provides information on their function for their host [50].

Denitrifying endosymbionts are proposed to perform respiration and generate ATP for their host, and were first thought to be strict anaerobes [16]. It was recently established that some groundwater species are in fact facultatively aerobic [17]. Ten out of our fourteen newly described wastewater symbiont species appear facultatively aerobic. Interestingly, the cytochrome-*cbb_3_* oxidase, a high-affinity terminal oxidase encoded in the symbiont genomes, was differentially lost at least two times among the symbiont lineages (Supplementary Discussion), most notably in the whole monophyletic clade including BJNG_1, ESTL_1 and *Ca*. A. michiganensis from wastewater, as well as the lacustrine *Ca*. A. ciliaticola. Thus, it is likely that the symbiont transition to an obligately anaerobic lifestyle occurred multiple times.

Some denitrifying symbionts and their hosts exist in oxic environments even without a terminal oxidase. For example, *Ca*. A. parvus, which lacks the cytochrome-*cbb_3_* oxidase was present in the aeration tank of the Seehausen WWTP. However, abundant anoxic microniches are likely present in these aeration tanks due to the typically low efficiency of aeration, use of aggregated sludge with high organic carbon content, and high aerobic respiration rates. Plagiopylean ciliates are thought to represent an anaerobic protist clade [51] and host detoxification mechanisms may also contribute to their survival under oxic conditions [52].

Due to its higher level of genome reduction, *Ca.* A. parvus appears to also lack other genes that are consistently conserved in other symbiont genomes, such as the last leftover genes of the TCA cycle (2-oxoacid:ferredoxin oxidoreductase and succinate-CoA ligase) and some aminoacyl-tRNA synthetases (*trpS*, *glyQ*, *glyS*, *thrS*). Notably, *Ca*. A. parvus is the first denitrifying symbiont to be missing a part of the denitrification pathway, namely the whole *nosDLFRYZ* operon, including genes for catalytic as well as maturation subunits. The transfer of *nosZ* and the accessory genes to the host genome or their presence on a plasmid as shown in *Methylocystis* sp. SC2 [53] appear unlikely as almost no read closely related to the *Azoamicaceae nosZ* sequences could be retrieved from the source metagenome of *Ca*. A. parvus.

While all symbiont genomes known to date encode a complete denitrification, this pathway is often modular among free-living denitrifiers [54]. It is thus possible that the gene complement for denitrification in symbiotic denitrifiers might be more flexible than what is currently known. As nitrous oxide reductase is the only known enzyme that can remove nitrous oxide [55], it is likely that N_2_O is the end product of denitrification in the globally distributed *Ca*. A. parvus.

Nitrous oxide is known to be produced in WWTPs due to incomplete denitrification [56], although the global nitrous oxide emission estimates from WWTPs remain uncertain [57,58]. Seasonal variations in nitrous oxide concentrations were detected [59] and associated with possible prokaryotic species in the WWTPs [60]. Our data point out that, although their relative abundance is likely quite lower than that of free-living *nosZ*-less denitrifiers, host-associated microbes in principle may be implicated in the production of nitrous oxide.

<u>Contribution of ciliates associated with denitrifying endosymbionts to wastewater treatment</u> Based on 18S rRNA gene identity, the hosts of the newly described denitrifying endosymbionts appear to belong to Plagiopylea and are closely related to previously reported hosts of lacustrine and groundwater symbionts [16,17]. In the Seehausen WWTP, different morphologies of the putative ciliate hosts were observed, which could indicate that the different wastewater symbionts associate with distinct Plagiopylean hosts. Alternatively, the delineation of this ciliate group may be better supported by molecular data, such as SSU rRNA gene sequences, rather than morphological criteria, as reported before for Plagiopylea [61] and other groups of anaerobic ciliates [62].

Ciliates have previously been found in distinct stages of wastewater treatment [63] and are common eukaryotes in WWTP. Indeed, Alveolata, to which ciliates belong, are after Fungi the second most abundant group of microeukaryotes in anaerobic WWTPs [64]. Their main role is thought to be shaping the prokaryotic community through predation, but the full involvement of ciliates in WWTP processes remains unclear [10,65]. For example, protist symbionts in WWTPs have been investigated for their potential pathogenicity [66] and contribution to methane production in anaerobic digesters [67]. This study shows that, despite their low abundance, protists and their symbionts may be active in biogeochemical processes and thus directly contribute to wastewater treatment rather than only via predation.

## Conclusions

Our study of global wastewater metagenomes and amplicons reveals that denitrifying endosymbionts are omnipresent in a variety of engineered and natural aquatic environments. We show that denitrifying endosymbionts and their ciliate hosts are common in WWTPs across the world, display strong temporal variability, and can survive in dynamic environmental conditions. Some symbiont lineages have even been detected in lab-based bioreactors, suggesting bioreactor cultivation as a promising avenue to obtain laboratory enrichments or axenic cultures of the ciliate host, which would allow to investigate their physiology and interaction with symbionts.

## Acknowledgements

We thank Klaus Reichel and *hanseWasser* Bremen GmbH for allowing us to sample the WWTP in Seehausen, Bram Vekeman and Paloma Garrido Amador for help with sampling, and Daniela Tienken for technical assistance. We thank Boran Kartal for helpful discussions.

## Author contributions

L.N., D.R.S. and J.M. designed the study. L.N. and L.M.Z. performed sampling at the Seehausen WWTP and DNA/RNA extraction. C.M.S., L.L. and M.K.D.D. contributed metagenomic data. L.N., D.R.S. and B.J.W. carried out bioinformatics analyses. L.M.Z. conducted the fluorescence in situ hybridisation and microscopy. L.N., D.R.S. and J.M. wrote the manuscript with contributions from all co-authors.

## Etymology

### Description of ‘*Candidatus* Azoamicus mariagerensis’, sp. nov

‘Candidatus Azoamicus mariagerensis’ (ma.ri.a.ger.en’sis. N.L. masc. adj. *mariagerensis,* originating from Mariager as this species was first identified in a wastewater treatment plant near the Mariagerfjord in Denmark). The two strains recovered from WWTPs in Mariager (Denmark) and Chelsea (USA) are referred to as *Ca.* A. mariagerensis MARI and *Ca.* A. mariagerensis CHLS respectively.

### Description of ‘*Candidatus* Azoamicus parvus’, sp. nov

‘Candidatus Azoamicus parvus’ (par’vus. L. masc. adj. *parvus,* small, referring to its highly reduced genome). This species is represented by the genome from a WWTP in New Zealand.

### Description of ‘*Candidatus* Azoamicus michiganensis’, sp. nov

‘Candidatus Azoamicus michiganensis’ (mi.chi.gan.en’sis. N.L. masc. adj. *michiganensis,* originating from Michigan as this species was first identified in a bioreactor inoculated with water from a wastewater treatment plant in the city of Ann Arbor, Michigan). This species is represented by the genome from a bioreactor inoculated with wastewater from the Ann Arbor WWTP (Michigan, USA).

## Supplementary material

Supplementary Figures and Tables S1-3 can be found in the Supplementary Material. Source files used to generate main and supplementary figures as well as Tables S4-8 will be available upon publication.

## Data availability

Metagenomic and metatranscriptomic data, endosymbiont MAGs obtained from short reads and host 18S rRNA gene sequences generated in this study have been deposited on NCBI and ENA. Accession numbers will be available upon publication.

