## Supplemental material for "Occurrence and temporal dynamics of denitrifying protist endosymbionts in the wastewater microbiome"

**Short title:** Denitrifying endosymbionts in wastewater

**Affiliations:**

[a]Max Planck Institute for Marine Microbiology, Celsiusstrasse 1, 28359 Bremen, Germany

[b]Division of Microbial Ecology, Centre for Microbiology and Environmental Systems Science, University of Vienna, Djerassiplatz 1, 1030 Vienna, Austria

[c]Centre for Microbiome Research, School of Biomedical Sciences, Queensland University of Technology, 37 Kent Street, Woolloongabba QLD 4102, Australia

[d]Center for Microbial Communities, Department of Chemistry and Bioscience, Aalborg University, Fredrik Bajers Vej 7K, 9220 Aalborg, Denmark

***Correspondence:**

Louison Nicolas-Asselineau

Max Planck Institute for Marine Microbiology, Celsiusstrasse 1, 28359 Bremen, Germany

**Table of content**

Supplementary figures 3

Supplementary tables 14

Supplementary methods 18

Read-based selection of metagenomes of interest for the recovery of denitrifying endosymbiont MAGs 18
MAG recovery from Nanopore long-read sequencing 18
Genome refining 19
Genome annotation 19
Probe design and evaluation for fluorescence *in situ* hybridisation 20
DNA and RNA extraction 21
Metagenome and metatranscriptome sequencing 22
Read mapping on the recovered genomes 22
Recovery of putative host 18S rRNA gene sequences 23
16S rRNA gene amplicon processing 23
Supplementary discussion 24
Description of the wastewater MAGs 24
Genera and species delineation 24
Occurrence of genes coding for energy production in the MAGs 25
Fluctuations of denitrifying endosymbiont abundances over time 26
Diversity of denitrifying endosymbionts in the Seehausen WWTP 27
References 28

### Supplementary figures

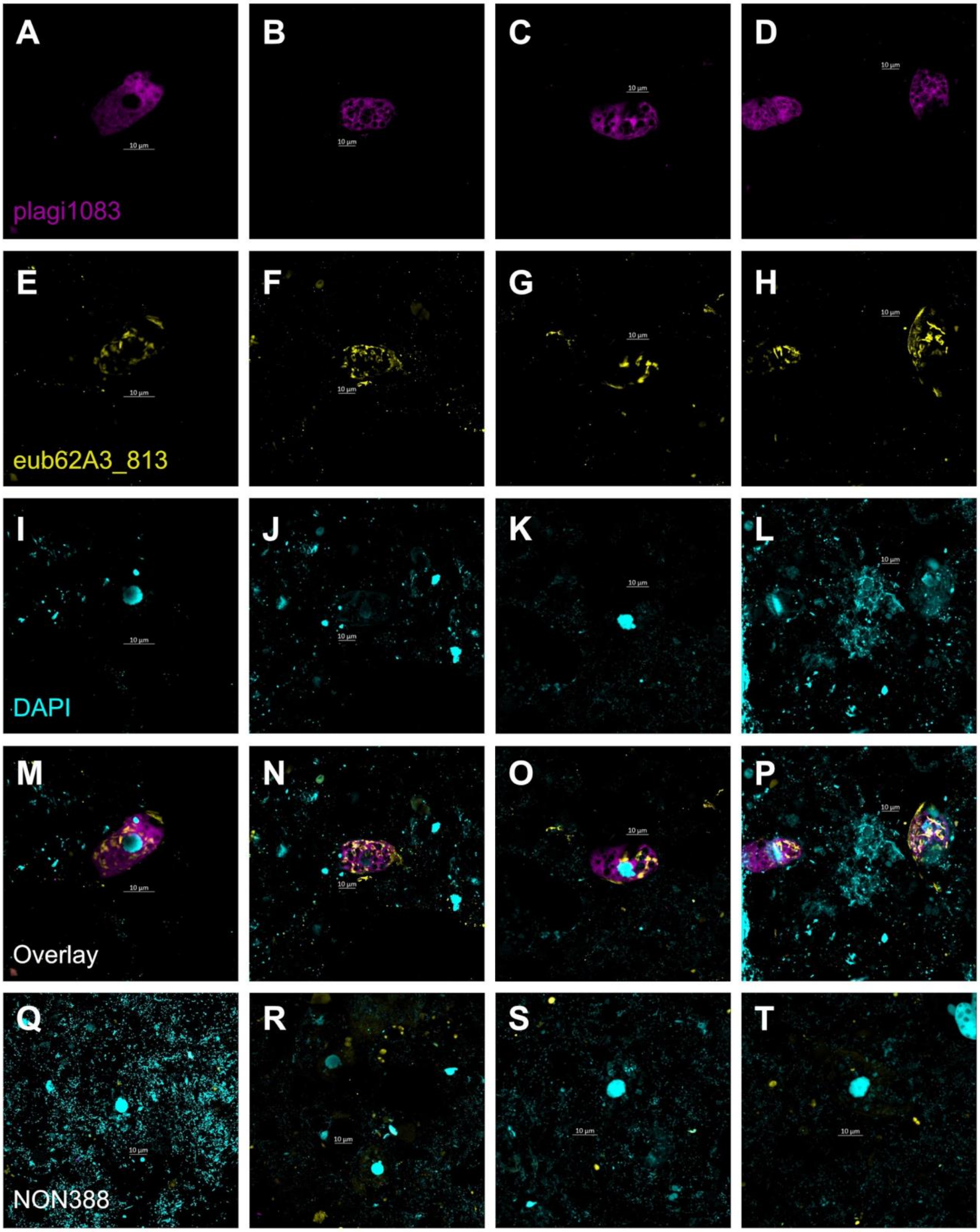

**Figure S1: Confocal laser scanning microscopy images of Plagiopylean ciliates from the Seehausen WWTP in Bremen, Germany after double labelling of oligonucleotide probes for FISH with A-D,** the plagi1083 probe targeting Plagiopylean ciliates, **E-H,** the eub62A3_813 probe targeting some *Azoamicus* endosymbionts, **I-L,** DAPI staining. **M-P,** Overlay of a-l images. **Q-T,** Negative control images with the NON388 probe.

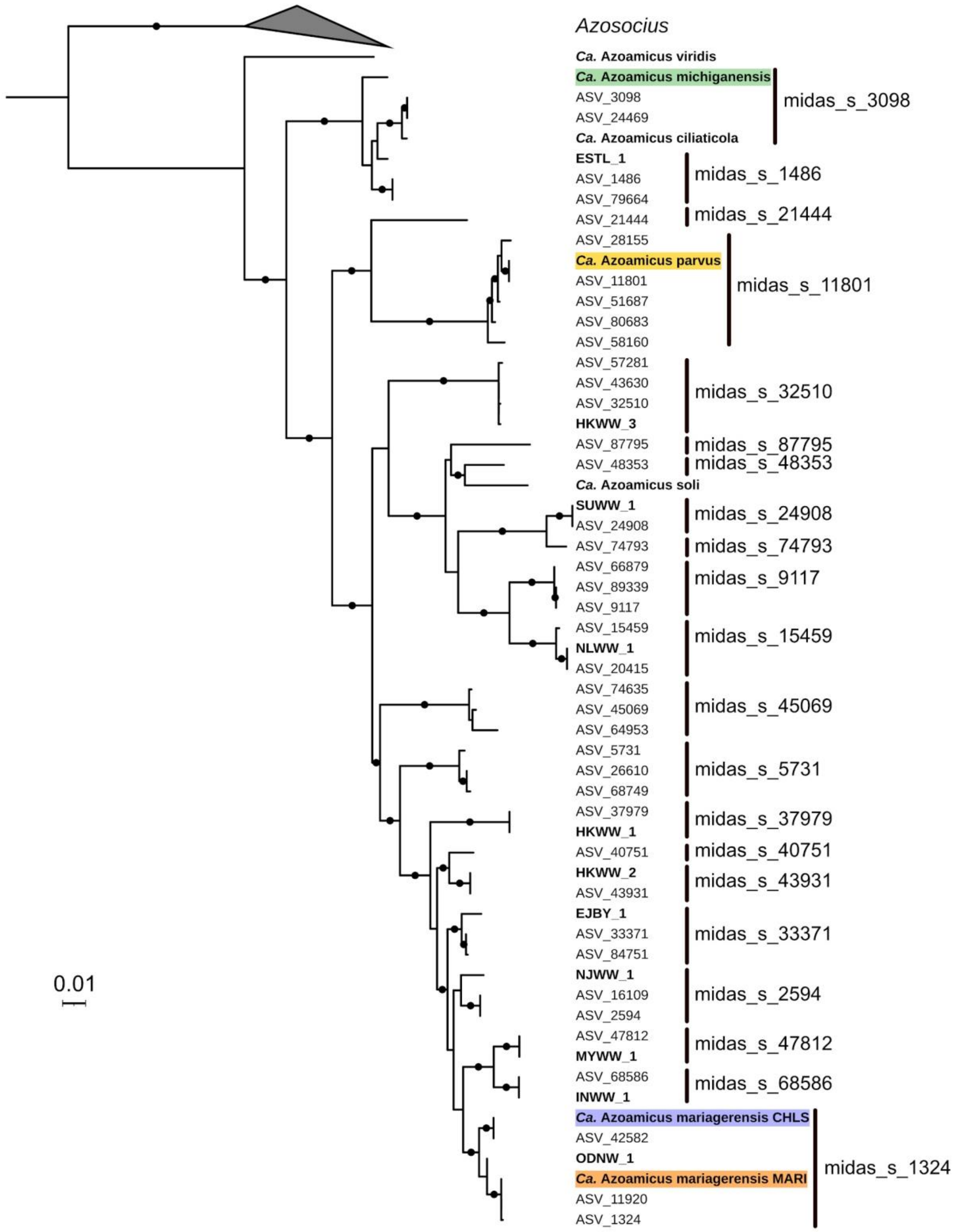


**Figure S2: 16S rRNA based approximate maximum likelihood phylogenetic tree of all putative *Azoamicaceae* members identified in the MiDAS 5.3 database and wastewater MAGs.** MAGs retrieved from wastewater are indicated in bold and complete genomes are highlighted. The

BJNG_1 MAG is not shown as no full-length 16S rRNA gene sequence could be recovered. Dots on branches indicate bootstrap values above 80%. The lineages from which a MAG was recovered are assigned to the MiDAS species with which they had the highest 16S rRNA gene sequence identity.

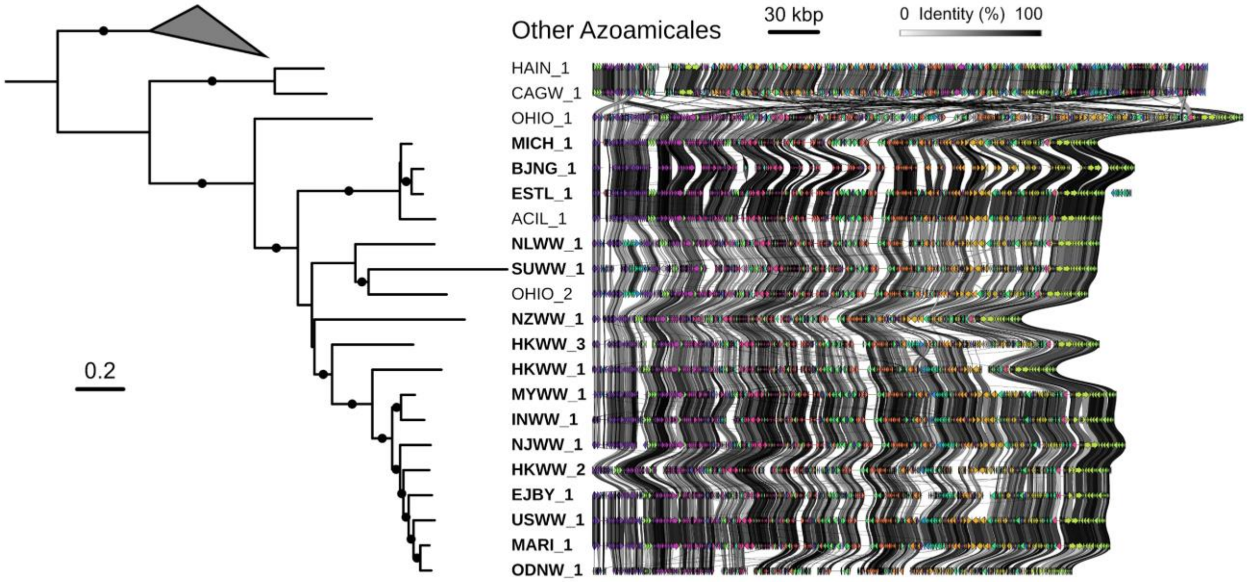


**Figure S3: Gene synteny in denitrifying endosymbionts.** Gene order conservation among the genomes of species belonging to *Ca.* Azosocius and *Ca.* Azoamicus. The phylogenomic tree is identical to the one shown in Fig. 1. CAGW_1: *Ca.* Azosocius agrarius, HAIN_1: *Ca*. Azosocius aquiferis, OHIO_1: *Ca.* Azoamicus viridis, MICH_1: *Ca.* Azoamicus michiganensis, ACIL_1: *Ca.* Azoamicus ciliaticola, NZWW_1: *Ca.* Azoamicus parvus, OHIO_2: *Ca.* Azoamicus soli, USWW_1: *Ca.* Azoamicus mariagerensis CHLS, MARI_1: *Ca.* Azoamicus mariagerensis MARI.

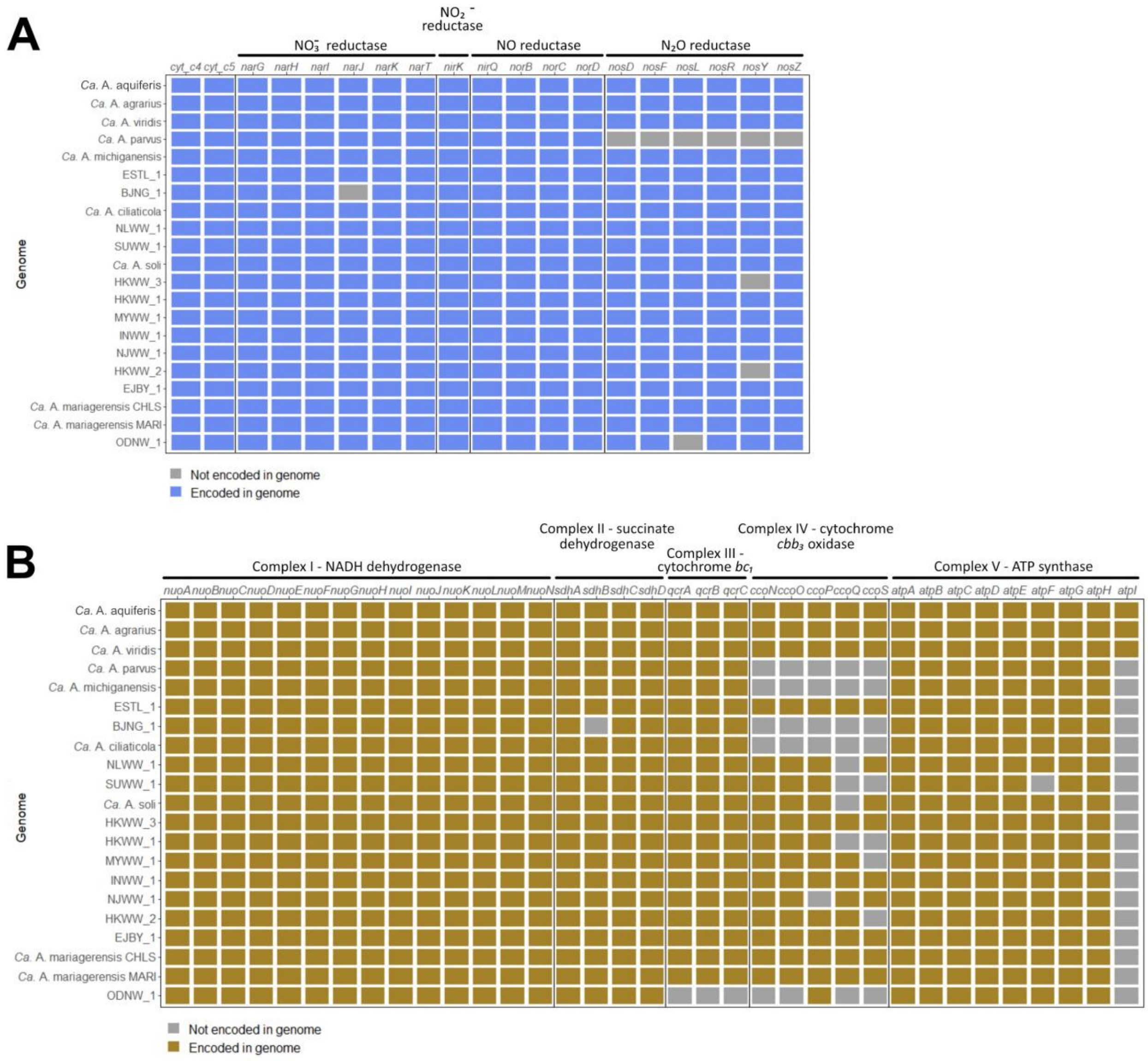


**Figure S4: Presence of genes coding for enzymes involved in energy production in the MAGs of denitrifying endosymbionts retrieved from wastewater.** Catalytic and non catalytic subunits of the complexes as well as accessory proteins are included. **A,** Genes coding for enzymes involved in denitrification. **B,** Genes encoding components of the electron transport chain.

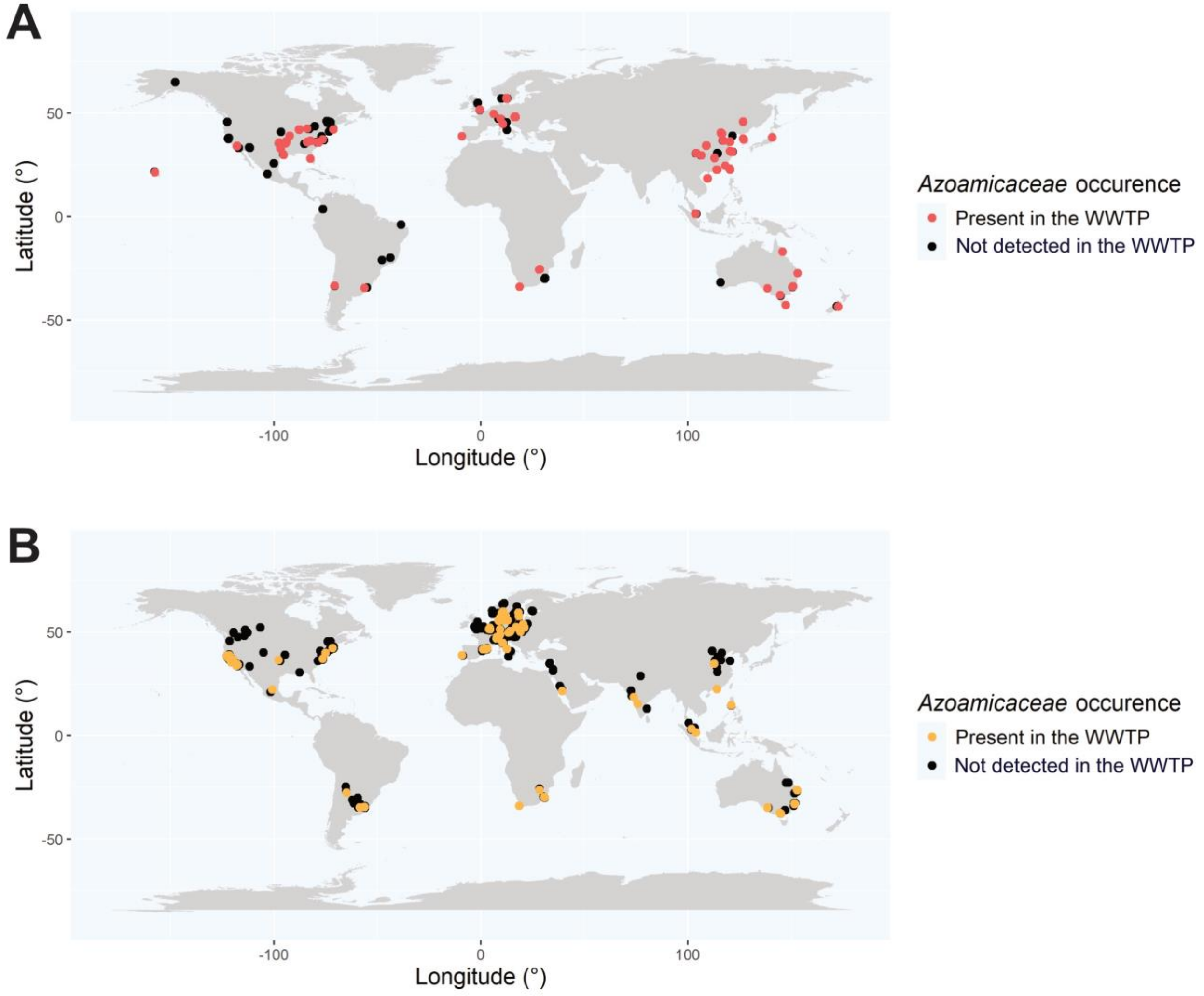


**Figure S5: Occurrence of *Azoamicaceae* in WWTPs sampled as part of the A, GWMC and B, MiDAS 4 campaigns**. Each dot corresponds to the location of one sampled WWTP with colours denoting the detection of *Azoamicaceae*.

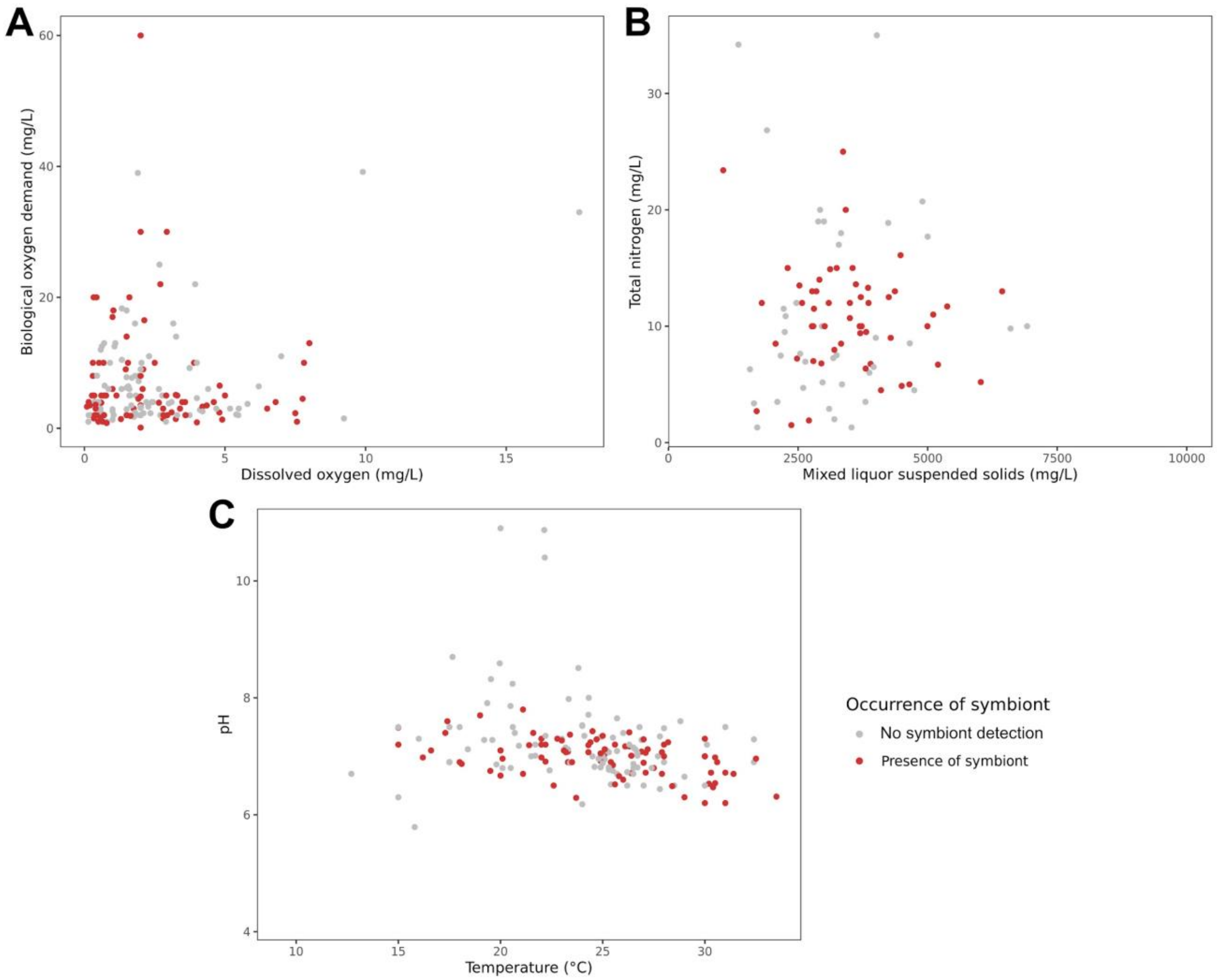


**Figure S6: Occurrence of denitrifying endosymbionts in GWMC 16S rRNA gene amplicons and the respective WWTP operating conditions.** Presence of symbionts was based on the detection of members of the midas_f_1324 family in the amplicon dataset. The WWTP parameters appear as presented in the source study metadata. **A,** Biological oxygen demand (mg/L) versus dissolved oxygen (mg/L). **B,** Total nitrogen (mg/L) versus mixed liquor suspended solids (mg/L). **C,** pH versus temperature (°C).

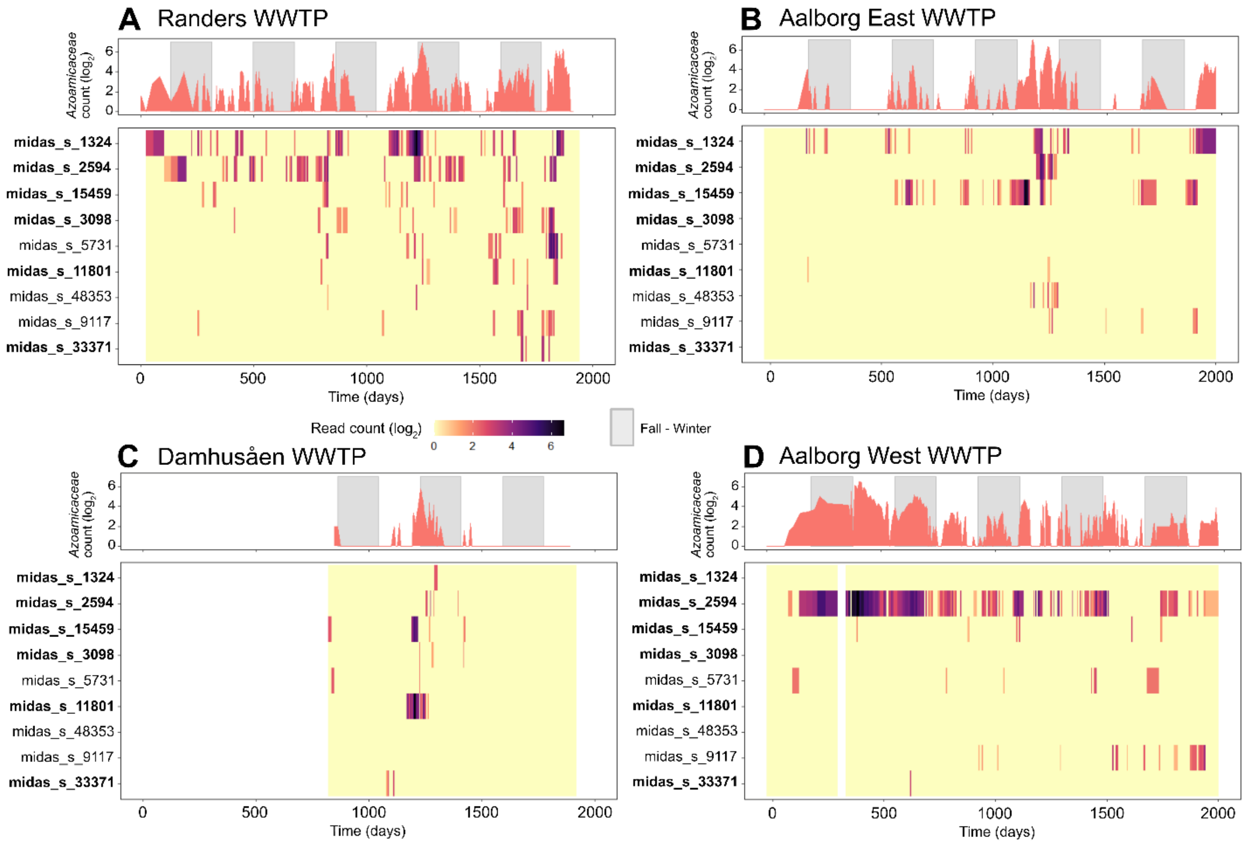


**Figure S7: Variations in denitrifying endosymbiont community composition and abundance over time in four Danish WWTPs.** Each panel depicts the community fluctuations at the family and species level in the **A,** Randers, **B,** Aalborg East, **C,** Damhusåen, **D,** Aalborg West WWTPs. All listed MiDAS 5.3 species are expected to belong to the *Azoamicaceae* family. A MAG is available for the species in bold. The abundance of a species is approximated by its 16S rRNA gene amplicon read count. Grey bars on the upper plots highlight fall-winter time defined as the period between 21 September and 21 March.

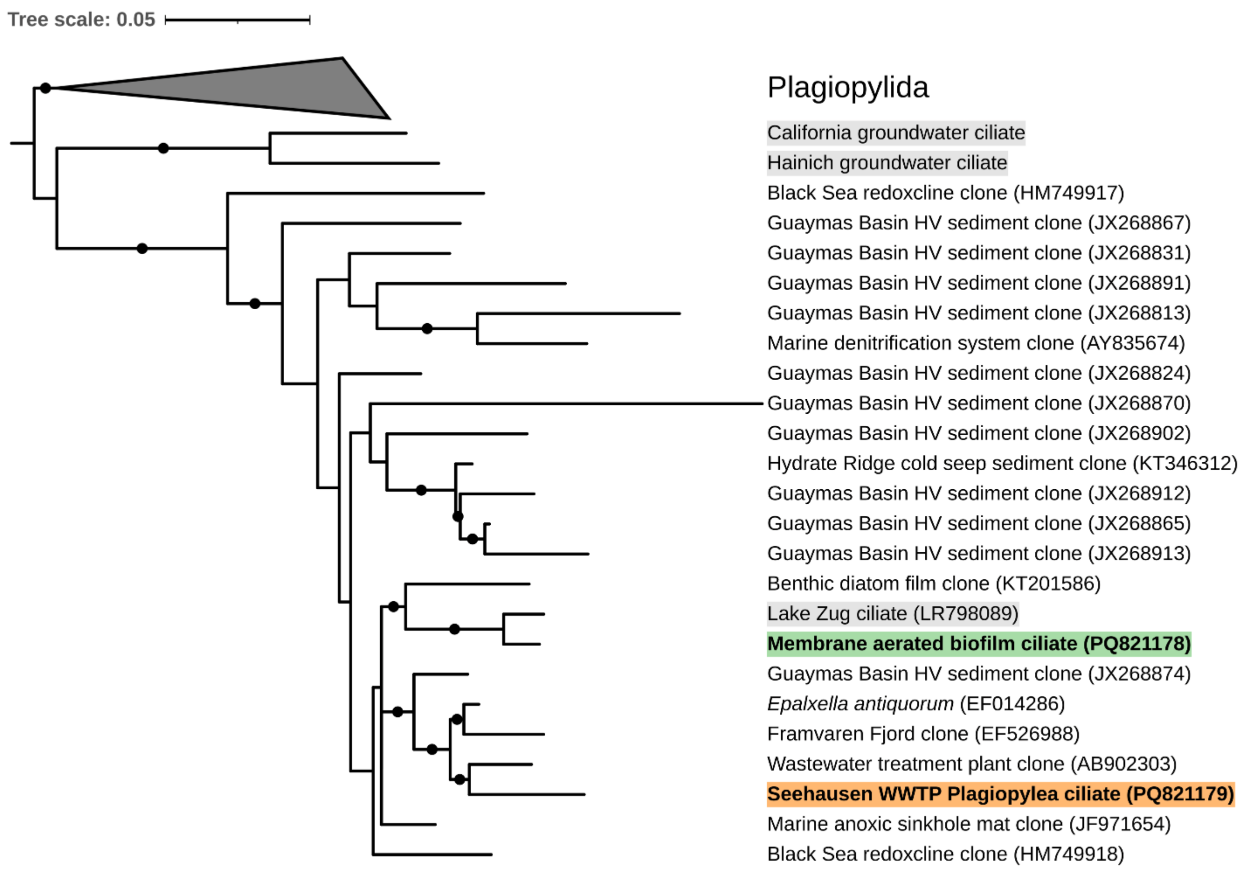


**Figure S8: 18S rRNA gene based maximum likelihood phylogenetic tree of putative denitrifying endosymbiont hosts.** The potential host of *Ca*. A. mariagerensis MARI from the Seehausen WWTP (orange background in bold) and *Ca*. A. michiganensis (green background in bold), as well as sequences of non-wastewater ciliates retrieved from datasets containing denitrifying endosymbionts (grey background) are included. Dots indicate bootstrap values above 80%.

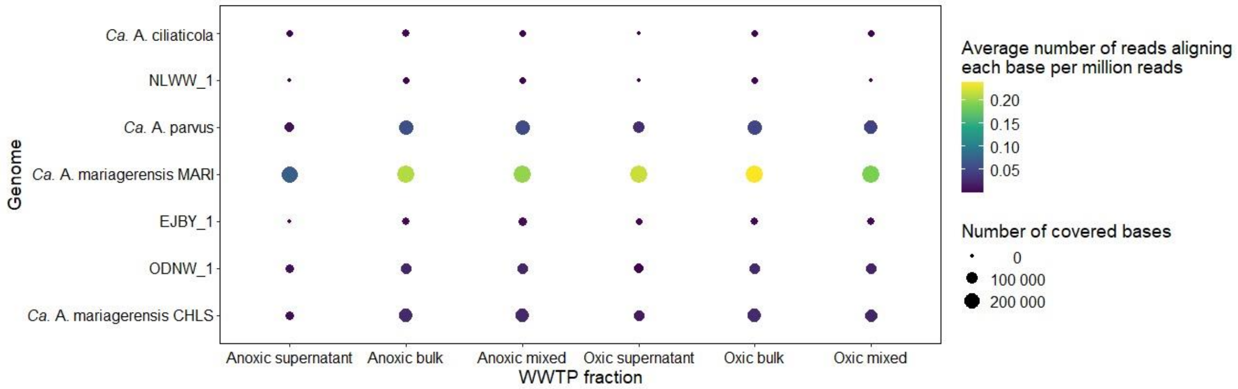


**Figure S9: Diversity of denitrifying endosymbiont lineages in WWTP fractions with various oxygen concentrations.** Abundance of denitrifying endosymbiont lineages in wastewater sampled in October 2022 in the aeration and anoxic tanks of the Seehausen WWTP in Bremen, Germany. The amount of reads mapping on each base is normalised per million reads in the metagenomic dataset.

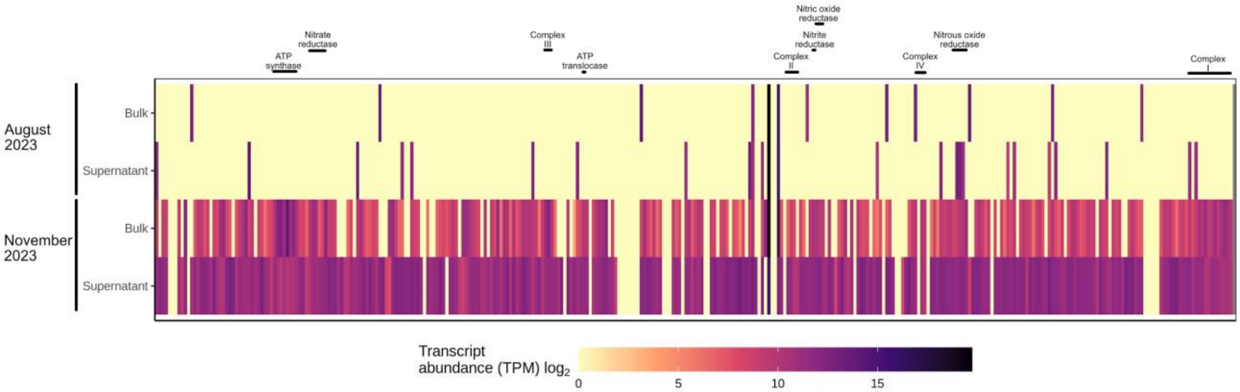


**Figure S10: Transcription level of genes of *Ca.* A. mariagerensis MARI in the Seehausen WWTP in Bremen, Germany, at two timepoints and in two wastewater fractions.** Each vertical bar corresponds to one gene and the count of mapping metatranscriptomic reads is normalised using the Transcripts Per Million method.

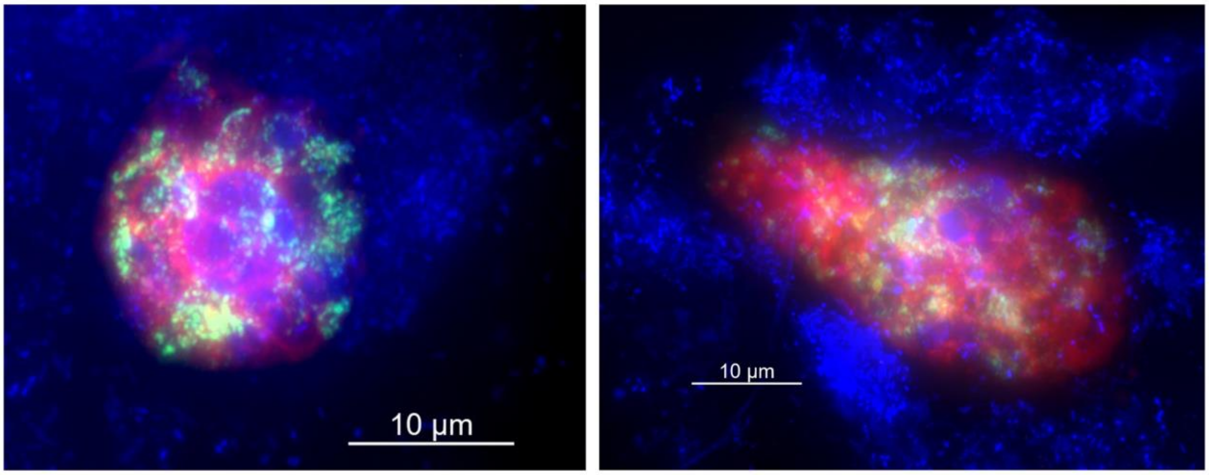


**Figure S11: Plagiopylean ciliates and denitrifying endosymbionts visualised by double labelling of oligonucleotide probes for FISH.** Images originate from the supernatant of the Seehausen WWTP in November 2023. The Plagiopylean probe is shown in red, the denitrifying endosymbiont specific probe in green and DAPI staining in blue. The visualised ciliates display different morphologies.

### Supplementary tables

**Table S1: Sampling of wastewater from the Seehausen WWTP, Bremen, Germany.** Bottles containing fresh samples were left undisturbed until the solid fraction had settled to the bottom and a clear supernatant fraction was visible on top. These fractions are respectively named “sludge” and “supernatant”.

| Sampling date | WWTP tank | Sampled fraction | Purpose | Volume (mL) |
|---|---|---|---|---|
| 17 October 2022 | Aeration tank | Supernatant | Metagenome sequencing | 375 |
| 17 October 2022 | Aeration tank | Sludge | Metagenome sequencing | 3 |
| 17 October 2022 | Anoxic tank | Supernatant | Metagenome sequencing | 100 |
| 17 October 2022 | Anoxic tank | Sludge | Metagenome sequencing | 3 |
| 28 August 2023 | Aeration tank | Supernatant | Metagenome sequencing | 200 |
| 28 August 2023 | Aeration tank | Supernatant | Metatranscriptome sequencing | 200 |
| 28 August 2023 | Aeration tank | Sludge | Metagenome sequencing | 3 |
| 28 August 2023 | Aeration tank | Sludge | Metatranscriptome sequencing | 10 |
| 14 November 2023 | Aeration tank | Supernatant | Metagenome sequencing | 100 |
| 14 November 2023 | Aeration tank | Supernatant | Metatranscriptome sequencing | 100 |
| 14 November 2023 | Aeration tank | Sludge | Metagenome sequencing | 3 |
| 14 November 2023 | Aeration tank | Sludge | Metatranscriptome sequencing | 10 |

### Table S2: FISH probes

| Probe name | Specificity | Sequence | Reference | Dye |
|---|---|---|---|---|
| eub62A3_813 | Most *Ca.* Azoamicus species | 5'-CTAACAGCAAGTTTTCATCGTTTA-3' | Graf et al., 2021 [1] | Atto488 |
| plagi1083 | Plagiopylean class with a weak mismatch to most *Trimyema* sequences | 5'-TTGTGTCCATACTTCCCCC-3' | This study | Atto594 |
| plagi1083 competitor 1 | | 5'-TTGCGACCATACTCCCCCC-3' | This study | |
| plagi1083 competitor 2 | | 5'-TTGCAACCATACTTCCCCC-3' | This study | |
| plagi1083 competitor 3 | | 5'-TTGTGTCCATACTACCCCC-3' | This study | |
| NON338 | | 5'-ACTCCTACGGGAGGCAGC-3' | Wallner et al., 1993 [2] | Atto488 |

**Table S3: Sequencing datasets.** Bottles containing fresh samples were left undisturbed until the solid fraction had settled to the bottom and a clear supernatant fraction was visible on top. These fractions are respectively named “sludge” and “supernatant”. A mixed fraction was taken after homogenising the two fractions in the sample.

| ID | Type | Sequencing technology | No. of (paired-end) reads | Total sequenced (Gb) | Sample description |
|---|---|---|---|---|---|
| MG_AN_SUP_2210 | DNA | NextSeq2000 (2x150 bp) | 35,636,650 | 10.7 | Supernatant wastewater, anoxic reactor, October 2022 |
| MG_AN_SLU_2210 | DNA | NextSeq2000 (2x150 bp) | 35,419,732 | 10.6 | Sludge wastewater, anoxic reactor, October 2022 |
| MG_AN_MIX_2210 | DNA | NextSeq2000 (2x150 bp) | 33,618,196 | 10.1 | Mixed wastewater, anoxic reactor, October 2022 |
| MG_OX_SUP_2210 | DNA | NextSeq2000 (2x150 bp) | 25,397,410 | 7.6 | Supernatant wastewater, oxic reactor, October 2022 |
| MG_OX_SLU_2210 | DNA | NextSeq2000 (2x150 bp) | 28,981,611 | 8.7 | Sludge wastewater, oxic reactor, October 2022 |
| MG_OX_MIX_2210 | DNA | NextSeq2000 (2x150 bp) | 28,571,024 | 8.6 | Mixed wastewater, oxic reactor, October 2022 |
| MG_OX_SUP_2308 | DNA | NextSeq2000 (2x150 bp) | 41,161,150 | 12.3 | Supernatant wastewater, oxic reactor, August 2023 |
| MG_OX_SLU_2308 | DNA | NextSeq2000 (2x150 bp) | 41,678,240 | 12.5 | Sludge wastewater, oxic reactor, August 2023 |
| MG_OX_SUP_2311 | DNA | NextSeq2000 (2x150 bp) | 40,464,411 | 12.1 | Supernatant wastewater, oxic reactor, November 2023 |
| MG_OX_SLU_2311 | DNA | NextSeq2000 (2x150 bp) | 38,493,773 | 11.5 | Sludge wastewater, oxic reactor, November 2023 |
| MT_OX_SUP_2308 | RNA | NextSeq2000 (2x150 bp) | 56,516,662 | 17.0 | Supernatant wastewater, oxic reactor, August 2023 |
| MT_OX_SLU_2 | RNA | NextSeq2000 | 56,978,760 | 17.1 | Sludge wastewater, oxic |

| | | | | | |
|---|---|---|---|---|---|
| 308 | | (2x150 bp) | | | reactor, August 2023 |
| MT_OX_SUP_2 311 | RNA | NextSeq2000 (2x150 bp) | 54,580,065 | 16.4 | Supernatant wastewater, oxic reactor, November 2023 |
| MT_OX_SLU_2 311 | RNA | NextSeq2000 (2x150 bp) | 56,103,661 | 16.8 | Sludge wastewater, oxic reactor, November 2023 |

**Table S4: Publicly available metagenomes screened for denitrifying endosymbiont sequences.** The screening consisted in identifying metagenomes of interest by detecting the presence of *tlcA* genes using the BLAST Score Ratio approach [3,4]. The accession of all screened metagenomes is indicated as well as the resource (GWMC/Sandpiper) used for the selection of this dataset. All whole genome sequencing datasets published by the GWMC [5] were screened. Sandpiper [6] allowed to identify metagenomes that included sequences related to those of denitrifying endosymbionts that were also screened. From the screening results, promising metagenomes were assembled, and MAGs were retrieved in some cases. The reason for keeping a MAG for further analysis or not is detailed in the “Comment” column.

**Table S5: Summary and characteristics of the MAGs of denitrifying endosymbionts recovered from wastewater.** Some MAG statistics are listed with metadata regarding the sample from which they were recovered.

**Table S6: Average Nucleotide Identity (ANI) and Average Amino Acid Identity (AAI) of all known species of denitrifying endosymbionts.**

**Table S7: Annotation table of the denitrifying endosymbiont MAGs recovered from wastewater.** KEGG, NCBI COG20 and Pfam annotation are shown as well as a manual consensus. The gene position is indicated for the complete MAGs and a presence/absence matrix is shown for the incomplete ones.

**Table S8: Transcripts per million values of all *Ca*. Azoamicus mariagerensis MARI genes in the bulk and supernatant wastewater from the Seehausen WWTP, sampled in August and November 2023.** The gene annotation was computed with Prokka 1.14.6 [7].

**Supplementary methods**

<u>Read-based selection of metagenomes of interest for the recovery of denitrifying endosymbiont MAGs</u>

All protein sequences encoded by known *tlcA* genes on the one hand and *nosZ* genes on the other hand were extracted from the Genome Taxonomy Database (GTDB, [8]) and Genomes from Earth's Microbiome (GEM, [9]) to form comprehensive TlcA and NosZ protein databases. Selected metagenomes were screened for the presence of both genes using the BLAST Score Ratio [3,4]. According to this approach, the presence of the *tlcA* gene on a read is assessed by comparing the BLAST score of the translated gene sequence against our custom TlcA to the maximum possible BLAST score, and similarly for *nosZ*.

If the *tlcA* gene was present in more than twenty reads, the metagenomic dataset was selected for investigation. Otherwise, the BLAST Score Ratio values for the *nosZ* gene were computed and datasets with more than 20 reads carrying *nosZ* were also chosen for investigation. Others were discarded.

*nosZ* was chosen as a secondary marker gene for denitrifying endosymbionts in the event of *tlcA*-based screening failure. However, this secondary *nosZ* screening did not allow us to select any metagenome that was not already identified by the *tlcA* screening.

<u>MAG recovery from Nanopore long-read sequencing</u>

MAGs of denitrifying endosymbionts were retrieved from long-read sequencing datasets generated using the PromethION flowcell (R10.4.1) and the ligation sequencing kit (SQK-LSK114). Guppy (super-accurate mode) was used for basecalling, and genome recovery was performed with the mmlong2 pipeline (https://github.com/Serka-M/mmlong2). Briefly, contigs were assembled with metaFlye v. 2.9.0 [10], polished once with medaka v. 1.8.0 (https://github.com/nanoporetech/medaka), and filtered to remove eukaryotic contigs using

Tiara v. 1.0.3 [11]. Binning was conducted with multiple tools, including MetaBAT2 v. 2.12.1 [12], GraphMB v. 0.2.4 [13], and SemiBin v. 1.4 [14], with the best bins selected by dRep v. 3.4.2 [15].

Genome refining

The MAGs recovered from short reads using the reference-based binning approach were refined in order to increase their completeness and reduce the number of contigs. Illumina reads from the source dataset of a MAG trimmed with TrimGalore v. 0.6.7 (https://github.com/FelixKrueger/TrimGalore) were mapped on the corresponding genome with an identity of 98% over a minimum of 30% of the read using minimap v. 2.17 [16] and CoverM v. 0.6.1 [17]. All reads mapping to the genome using these parameters were subsequently re-assembled with SPAdes v. 3.15.3 [18] using k-mer sizes of 27, 37, 47, 57, 67, 77, 87, 97, 107, 117 and 127 resulting in a new set of contigs. This iterative assembly process was aimed at extending the existing contigs and repeated ten times in order to increase the contig lengths as much as possible. Only contigs with a GC content and a coverage in the expected range were kept to reduce the risk of contamination. Contigs shorter than 1000 bp on which no feature could be identified were discarded.

Genome annotation

The starting position of the MAGs was reset to the origin of replication based on the GC skew by aligning the MAGs with the previously recovered complete genomes of the *Azoamicus* genus [1,19] in progressiveMauve [20]. Open reading frames were identified in the re-oriented MAGs with Prodigal v. 2.6.3 [21]. Then, the genomes were annotated using KEGG [22], NCBI COGs [23] and Pfam [24]. Transfer RNAs were also detected with tRNAScan-SE [25]. To further refine the annotation, the gene content of the MAGs was compared with that of *Ca.* A. viridis by using the reciprocal best hit (RBH) approach with diamond described in [26]. After blasting all MAGs with each other using the getRBH.pl script (https://github.com/Computational-conSequences/SequenceTools), the RBH results were merged in order to remove duplicate hits and obtain a table with all identified genes and their position in the genomes when they were present. This annotation table was manually corrected to establish a consensus in the gene functions and verify the positions in the genome. To assess the similarity of these MAGs, their ANI and AAI were computed with FastANI v. 1.33 [27] and CompareM v. 0.1.2 (https://github.com/dparks1134/CompareM).

Probe design and evaluation for fluorescence *in situ* hybridisation

To determine the identity of potential hosts of denitrifying endosymbionts, a specific oligonucleotide probe was designed in ARB v. 6.1 [28], using the probe design tool, based on the Plagiopylean 18S rRNA gene sequences available in the SILVA SSU Ref NR 99 132 database [29]. The specific oligonucleotide probe plagi1083 (5'-TTGTGTCCATACTTCCCCC-3') targets most members of the Plagiopylean class but has a weak mismatch to most *Trimyema* sequences. The probe does not show non-target binding, but a range of sequences have only 1 or 2 mismatches to the probe sequence. To exclude non-target binding three competitor oligonucleotides, perfectly binding the majority of these sequences, were designed (competitor 1: 5'-TTGCGACCATACTCCCCCC-3'; competitor 2: 5'-TTGCAACCATACTTCCCCC-3'; competitor 3: 5'-TTGTGTCCATACTACCCCC-3').

To evaluate the specific formamide concentration, the double labelled oligonucleotide probe plagi1083 with dye Atto594 (Biomers, Germany) was used to label known Plagiopylean ciliates sampled in July 2021 in Lake Zug, Switzerland, at depths 175m and 190m. From each depth, 500 mL were sampled and fixed with 2% paraformaldehyde for 1-2h and filtered onto 3 µm-Isopore™ polycarbonate filters (Merck Millipore, USA). The FISH protocol is based on Glöckner et al., 1996 with the following changes: filters were cut into pieces and embedded in 0.2% MetaPhor™ Agarose (Lonza, Switzerland). The filters were submerged in hybridization buffer containing 0, 10, 20, 25, 30, 35, 40, 45, 50 or 70% formamide and 5ng probe $\mu L^{-1}$ (the same concentration of each competitor was supplied), and incubated for 2h at 46°C. Filters were washed first in pre-heated washing buffer (NaCl concentrations were adjusted accordingly [30]) for 10 minutes at 48°C, then in ice cold milliQ water and dried at room temperature. Filters were embedded in 4:1 Citifluor Vectashield (Vector Laboratories, USA) and observed under the Axio IMager M2 epifluorescence microscope (Zeiss, Germany) in 400 x magnification. The exposure time was adjusted, based on the signal intensity at 10% formamide, to 88 000 ms. Up to a formamide concentration of 35%, all the targeted ciliates seemed to have been labelled. From 40% formamide, only a fraction was hybridised. No labelling was visible above a formamide concentration of 50%. Non-target binding was not observed. The probe was thus further used with a formamide concentration of 35%.

Additionally, the previously described eub62A3_813 probe [1] designed to target *Ca*. Azoamicus symbionts was used for the visualisation of denitrifying endosymbionts. Since the groundwater and wastewater MAGs have expanded the known diversity of the *Ca*. Azoamicus genus, we first re-evaluated the specificity of this probe. While the probe still specifically targets *Ca*. Azoamicus, it does not cover all new species, notably *Ca.* A. parvus, that is the most widespread. Importantly, it does target *Ca*. A. mariagerensis MARI, which was typically the most abundant symbiont species in the Seehausen WWTP.

DNA and RNA extraction

Samples for DNA/RNA extraction were preserved within 2 hours of sampling. The respective samples are described in Table S1. Supernatant fractions were filtered onto a 0.22 µm-Sterivex filter™ (Merck Millipore, USA). The 3-10 mL of solid fractions were centrifuged at 4°C for 20 minutes at 7190 x g. The supernatant was discarded, and the pellet used for DNA/RNA extraction. The filters and pellets were either stored at -20°C for nucleic acid extraction within one day or frozen in liquid nitrogen and stored at -80°C for longer term storage. DNA and RNA were extracted simultaneously using the ZymoBIOMICS™ DNA/RNA MiniPrep Kit (Zymo Research, USA). The pellets were dissolved in 750µL DNA/RNA shield (Zymo Research, USA), and filters were cut into pieces and then transferred to 750µL DNA/RNA shield. Samples were transferred to BashingBead Lysis Tubes and vortexed at maximum speed for 30 minutes, followed by a 5 minute-centrifugation at 12 000 x g. DNA and RNA were extracted from this supernatant following the manufacturer's “DNA and RNA purification protocol”, including the DNase treatment for the RNA samples.

The DNA of the bulk fraction sampled in October 2022 was extracted using the DNeasy® PowerSoil®(QIAGEN, Netherlands) kit and the protocol provided by the manufacturer, with the addition that elution buffer C6 was preheated to 55°C before use. The RNA and supernatant samples from October 2022 were extracted with the ZymoBIOMICS™ DNA/RNA MiniPrep Kit as described above.

Metagenome and metatranscriptome sequencing

All DNA samples were fragmented and processed for an Illumina-compatible library with NEBNext Ultra™ II FS DNA Library Prep Kit for Illumina (New England Biolabs, USA). Total RNA was processed to generate an Illumina-compatible library with Universal Prokaryotic RNA-Seq Library Preparation Kit (Tecan Genomics, USA) including rRNA depletion.

For both DNA and RNA, sequencing-by-synthesis was performed on the Illumina NextSeq 2000 sequencer (Illumina Inc., USA) in 2 x 150 bp paired-end read mode. Library preparation, and sequencing were conducted by the Max Planck-Genome-Center Cologne, Germany (https://mpgc.mpipz.mpg.de/home/). A detailed description of the sequencing datasets can be found in Table S3.

Read mapping on the recovered genomes

The presence and diversity of denitrifying endosymbiont species in the Seehausen WWTP and their level of gene transcription were determined by mapping respectively the metagenomic and metatranscriptomic reads on the wastewater MAGs. Reads sequenced from the distinct wastewater fractions were trimmed as described in the main Material and Methods ("Recovery of denitrifying endosymbiont genomes" section). They were then recruited on the MAGs with an identity of 95% over a minimum of 80% of the read through the covered bases and mean method using CoverM v. 0.6.1.

The presence of a genome in a given metagenome was assessed using the previously determined relation between breadth, i.e. the fraction of a genome covered by at least one read, and depth of coverage, i.e. the mean number of reads aligning each base of a genome [31,32]. A genome was considered present in a metagenome when the calculated breadth of coverage was within 15% of the expected breadth according to the formula: expected breadth

= 1 - e^(-0.883*depth of coverage). The observed and expected values were plotted as a scatter plot and a line plot respectively using R v. 4.2.1.

The transcripts per million values of each gene were computed as described in [33] using the number of reads mapped on the individual protein-coding genes of the wastewater MAGs identified by Prokka v. 1.14.6 [7].

Recovery of putative host 18S rRNA gene sequences

For 18S rRNA based phylogenetic analyses of denitrifying endosymbiont hosts, the metagenomic datasets from which wastewater MAGs were retrieved were run in phyloFlash v. 3.4.2 [34]. One full-length 18S rRNA gene sequence similar to the one from the Lake Zug host could be obtained from datasets SRR10823758 and SRR10823760, presumably belonging to the host of *Ca.* A. michiganensis. Additionally, the unassembled reads from the Bremen WWTP metagenome classified as belonging to Plagiopylea by phyloFlash were selected and assembled with MEGAHIT v. 1.2.9 resulting in a 2 kbp-long contig. This contig was blasted against the NCBI nr database and subset to extract the sequence aligning with the top hit, *Epalxella antiquorum* 18S gene (accession EF014286.1). The resulting sequence supposedly belonged to a host of denitrifying endosymbionts.

16S rRNA gene amplicon processing

No rarefaction was done on the amplicon datasets before processing. The reads were first imported into QIIME 2 v. 2022.11.1 [35] and subsequently trimmed using the cutadapt tool [36] to remove the corresponding primers (27F and 534R for the MiDAS 4 paired-end reads and time series datasets, 515F and 806R for the MiDAS 4 single-end reads and GWMC reads). The reads were then truncated to keep a Phred score above 20 with the DADA2 package [37]. Representative sequences were retrieved by DADA2 and clustered into ASVs with a 99% identity cut-off using VSEARCH [38]. The MiDAS v. 5.3 reference sequences and taxonomy (https://www.midasfieldguide.org/guide/downloads) were imported into QIIME 2 and used to train a classifier with the feature-classifier [39] and Scikit-learn plugins [40]. The trained classifier assigned a taxonomy to the clustered ASVs using the two aforementioned plugins. The results for paired-end and single-end amplicons belonging to the same WWTP were eventually combined.

### Supplementary discussion

Description of the wastewater MAGs

Two MAGs retrieved from WWTPs in Mariager (Denmark) and Chelsea (USA) had a similar size and GC content of 297 143 bp (24.0% GC) and 294 646 bp (23.8% GC), respectively. Furthermore, based on their high 16S rRNA gene identity of 98.6% we consider these two MAGs as two strains of one species, which we name *Candidatus* Azoamicus mariagerensis sp. nov.

Another complete MAG, assembled from a WWTP near Christchurch in New Zealand, displayed a 16S rRNA gene that was 91.9% similar to that of *Ca.* A. mariagerensis MARI. While its general genome organization is comparable to that of *Ca.* A. mariagerensis, the GC skew pattern differs. Therefore, we propose that this MAG represents a new species, which we name *Candidatus* Azoamicus parvus, sp. nov. Interestingly, *Ca.* A. parvus possesses the most reduced genome of all denitrifying endosymbionts described so far (246 131 bp; 20.6 % GC), which is nearly 40 kbp smaller than the second shortest genome, that of *Ca*. A. soli (284 kpb). As a result, *Ca.* A. parvus genome encodes fewer genes (295 genes), and has a higher protein coding density (93%).

The last complete MAG is 289 828 bp long, with a GC content of 23.7%, and a 16S rRNA identity of 92.5% and 90.1% to *Ca*. A. mariagerensis MARI and *Ca.* A. parvus respectively. This MAG was retrieved from a membrane-aerated biofilm reactor inoculated with wastewater from the Ann Arbor WWTP in Michigan, USA. We propose to classify this MAG as a new species which we name *Candidatus* Azoamicus michiganensis, sp. nov.

Genera and species delineation

Previously described species of denitrifying endosymbionts were classified into two genera, *Ca.* Azoamicus and *Ca.* Azosocius, based on 16S rRNA gene identity, ANI, AAI and pronounced gene synteny within each genus [19]. As denitrifying endosymbionts are fast-evolving organisms, it is likely that sequence identity metrics typically used to distinguish bacterial species and genera

overestimate their phylogenetic distance. Conservation of genome structure was thus proposed as an additional criterion for genus designation [19]. Based on the gene synteny argument, all wastewater MAGs were assigned to the *Ca*. Azoamicus genus (Fig. S3).
Furthermore, species delineation for microorganisms with tiny genomes is also complex. ODNW_1 shares an ANI above the species cut-off of 95% [41] with *Ca.* A. mariagerensis CHLS (Table S6) and is thus considered to belong to the *Ca.* A. mariagerensis species. ANI values among the other wastewater MAGs range between 76 and 90%, which is quite lower than the published threshold. 16S rRNA gene identity is also always below species boundaries [42]. Consequently, we choose to classify the remaining 13 wastewater MAGs into 13 distinct new species.

Occurrence of genes coding for energy production in the MAGs

Denitrifying enzymes are encoded by operons consisting of genes coding for catalytic subunits as well as accessory proteins, such as transporters and maturation proteins. The NarK and NarT transporters as well as the NarJ, NosR, NosD, NosF, NosL and NosY enzyme maturation proteins are encoded in all complete denitrifying endosymbiont genomes described in this study, with the exception of that of *Ca.* A. parvus that is missing the entire *nos* operon (Fig. S4). In almost all denitrifying symbionts, these enzymes form a complete functional denitrification chain.
The potential to respire oxygen is determined by the presence of genes coding for a terminal oxidase. Denitrifying symbiont genomes of lineages identified in groundwater were shown to possess a *cco* operon coding for a high affinity cytochrome-$cbb_3$ oxidase in their genome [19], revealing that these symbionts and their hosts may use oxygen in addition to nitrate as an electron acceptor. Similarly, most here-retrieved MAGs also include genes coding for a cytochrome-$cbb_3$ oxidase.
The *ccoNOPQ* operon is absent in some of the incomplete MAGs. We speculate that the MAG of ODNW_1 likely also includes a cytochrome-$cbb_3$ oxidase, as indicated by its phylogenomic placement (Fig. 1b) and by the presence of a subset of genes coding for complex IV (Fig. S4).
A *cco* operon of *Azoamicaceae* origin and coding for a potentially complete complex IV was retrieved from the source metagenome of the ESTL_1 lineage. Yet, this operon is located on its

own in a separate contig and not at the locus of complex IV that would be expected based on gene synteny in the *Azoamicus* genus. Therefore, there may be other denitrifying endosymbionts in the corresponding sample that remained undetected. Based on the genomic similarity of ESTL_1 with other species of symbionts that do not have a cytochrome-$cbb_3$ oxidase encoded in their genome (Fig. 1b), we chose to exclude the *cco* operon from the ESTL_1 MAG. Finally, like *Ca*. A. soli, HKWW_1 and SUWW_1 appear to have lost the *ccoQ* gene coding for a non-catalytic subunit but still encode the *ccoNOP* genes (Fig. S4).

Interestingly, both the *cco* and *nos* operons are missing from *Ca*. A. parvus. Although these operons are located close to each other in *Ca.* Azoamicus genomes, the genes located in between the two loci are still present in the MAG of *Ca*. A. parvus. This observation indicates that the two operons got independently lost in this species.

Due to its heavily reduced genome, *Ca.* A. parvus has lost more genes compared to other denitrifying endosymbionts. First, the genes coding for 2-oxoacid:ferredoxin oxidoreductase and succinate-CoA ligase are absent. These were the only genes dedicated to the TCA cycle left in the genome of *Ca.* A. ciliaticola [1] and *Ca.* A. parvus does not appear to have substitute genes for it. Genes coding for malate dehydrogenase and succinate dehydrogenase involved in the TCA cycle are still found in the *Ca.* A. parvus genome, but they may be part of other metabolic pathways [43,44]. Alternatively, the host could provide the required enzymes for a TCA cycle.

Fluctuations of denitrifying endosymbiont abundances over time

The relative abundance of denitrifying symbionts in the four assessed Danish WWTPs typically did not exceed 0.37% of the reads in the time-series amplicons, and these organisms do not seem to belong to the wastewater loose core taxa (core ASVs defined as top 0.1% most abundant ASVs in > 20% of samples, [45]). Only a tiny portion of ASVs identified in WWTPs sampled by the MiDAS 4 are in fact part of the core bacterial community [45]. Nevertheless, the midas_g_1324 genus that includes 10 species of denitrifying endosymbionts identified by MiDAS can be considered a conditionally rare or abundant taxon [45], meaning that its relative abundance was typically low but could occasionally increase to above 1% in a given WWTP.

Among the four investigated WWTPs, the Damhusåen WWTP displayed a somewhat different temporal abundance pattern. Denitrifying endosymbionts were not detected at all during most of the recorded time series, except for one large peak in 2018. This peak was due to a bloom of midas_s_11801 (*Ca*. A. parvus), a species that was very rare in the other three WWTPs.

Diversity of denitrifying endosymbionts in the Seehausen WWTP

The abundance of denitrifying endosymbiont lineages in the various samples taken from the Seehausen WWTP was estimated (Fig. 4, Fig. S9). The predicted relation between the breadth and depth of coverage of a genome in the metagenome was used to detect the true presence of a specific reference genome in the WWTP. In the obtained metagenomes of comparable sequencing depth, multiple species of denitrifying symbionts coexist in the reactors with uneven abundances, yet the community size and composition fluctuate with time as observed in the Danish WWTPs. *Ca*. A. mariagerensis MARI is present in high abundance in samples collected around fall (October 2022, November 2023) compared to other species, as its coverage is 6-7 fold higher than the second most abundant denitrifying endosymbiont lineage, *Ca.* A. parvus. *Ca*. A. ciliaticola, originally discovered in the anoxic layer of a freshwater lake and lacking a terminal oxidase, was detected in low abundance in the aeration tank in November 2023. The detection of ODNW_1 was less clear than that of other symbiont lineages found in the plant, which could be due to microdiversity within the *Ca*. A. mariagerensis species and unspecific read recruitment. Interestingly, denitrifying endosymbionts were found both in the supernatant and in the solid fraction of wastewater but consistently with a higher abundance in the latter. Based on these observations the symbionts might prefer the sludge fraction of wastewater, suggesting that they may be particle-associated.

Intriguingly, the abundance of *Ca.* A. parvus was relatively high at the three sampling points in the Seehausen WWTP. *Ca*. A. parvus is the most prevalent denitrifying endosymbiont lineage in WWTPs worldwide according to our global survey. One hypothesis explaining this observation would be that this species is well adapted to the dynamic conditions of WWTPs and consumes little energy due to its low amount of genes. Thus, the time of sampling would be less important than for other species and *Ca*. A. parvus would thus be detected more often.

Contrary to this assumption, *Ca*. A. parvus was mostly undetectable in the time series amplicons from the four WWTPs in Denmark (Fig. 3), making the ecological lifestyle of this species unclear.

Finally, metatranscriptomic data collected from different fractions of the Seehausen WWTP revealed that transcripts belonging to *Ca*. A. mariagerensis MARI are consistent with the abundance of the symbiont species in that plant. The vast majority of genes were transcribed both in the supernatant and solid fractions sampled in November 2023 (Fig. S10). Transcripts were detectable for genes involved in denitrification, electron transport chain and ATP exchange, suggesting that these pathways were functional. In particular, genes coding for subunits of the cytochrome-$cbb_3$ oxidase are transcribed, which is coherent with the fact that these samples were taken from an oxic environment. Moreover, *Ca*. A. parvus that possesses a smaller amount of genes than the other described denitrifying endosymbionts appears to only transcribe a few of them reducing its metabolic capacities even more (data not shown). The *nar*, *nir* and *nor* genes are transcribed for the denitrification pathway, and nitrous oxide might thus be produced.

38. Rognes T, Flouri T, Nichols B *et al.* VSEARCH: a versatile open source tool for metagenomics.

*PeerJ* 2016;**4**:e2584.
